# Methyltransferase SMYD5 Exaggerates IBD by Downregulating Mitochondrial Functions via Post-translational Control of PGC-1α Stability

**DOI:** 10.1101/2020.11.16.385765

**Authors:** Yuning Hou, Xiaonan Sun, Pooneh Tavakoley Gheinani, Xiaoqing Guan, Shaligram Sharma, Yu Zhou, Chengliu Jin, Zhe Yang, Anjaparavanda P. Naren, Jun Yin, Timothy L. Denning, Andrew T. Gewirtz, Zhonglin Xie, Chunying Li

**Affiliations:** Center for Molecular and Translational Medicine, Georgia State University, Atlanta, GA 30303; Division of Vascular Surgery, The First Affiliated Hospital, Sun Yat-Sen University, Guangzhou, China; Transgenic and Gene Targeting Core, Georgia State University, Atlanta, GA 30303; Department of Biochemistry, Microbiology, and Immunology, Wayne State University School of Medicine, Detroit, MI 48201; Department of Pediatrics, Division of Pulmonary Medicine, Cincinnati Children’s Hospital Medical Center, Cincinnati, OH 45229; Department of Chemistry, Center for Diagnostics & Therapeutics, Georgia State University, Atlanta, GA 30303; Center for Inflammation, Immunity and Infection, Institute for Biomedical Sciences, Georgia State University, Atlanta, GA 30303

**Keywords:** IBD, colitis, SMYD5, PGC-1α, Mitochondrion

## Abstract

**Background and Aims:** The expression and role of methyltransferase SET and MYND domain-containing protein 5 (SMYD5) in inflammatory bowel diseases (IBD) is completely unknown. Here, we investigated the role and the underlying mechanism of epithelial SMYD5 in IBD pathogenesis and progression.

**Methods:** The expression and subcellular localization of SMYD5 and peroxisome proliferator-activated receptor gamma coactivator-1α (PGC-1α) were examined by Western blot analysis, immunofluorescence staining, and immunohistochemistry in intestinal epithelial cells (IECs) and in colon tissues from human IBD patients and mice with experimental colitis. Mice with Smyd5 conditional knockout in IECs and littermate controls were subjected to DSS-induced experimental colitis and the disease severity and inflammation were assessed. SMYD5-regulated mitochondrial biogenesis was examined by RT-qPCR and transmission electron microscopy and mitochondrial oxygen consumption rate was measured in a Seahorse Analyzer system. The interaction between SMYD5 and PGC-1α was determined by co-immunoprecipitation assay. PGC-1α degradation and turnover (half-life) were analyzed by cycloheximide chase assay. SMYD5-mediated PGC-1α methylation was measured via *in vitro* methylation followed by mass spectrometry to identify the specific lysine residues that were methylated.

**Results:** Up-regulated SMYD5 and down-regulated PGC-1α were observed in IECs from IBD patients and mice with experimental colitis. However, Smyd5 depletion in IECs protected mice from DSS-induced colitis. SMYD5 was critically involved in regulating mitochondrial biology such as mitochondrial biogenesis, respiration, and apoptosis. Mechanistically, SMYD5 regulated mitochondrial functions in a PGC-1α dependent manner. Further, SMYD5 mediated lysine methylation of PGC-1α and facilitated its ubiquitination and proteasomal degradation.

**Conclusion:** SMYD5 attenuates mitochondrial functions in IECs and promotes IBD progression by enhancing the proteasome-mediated degradation of PGC-1α protein in methylation-dependent manner. Strategies to decrease SMYD5 expression and/or increase PGC-1α expression in IECs might be a promising therapeutic approach to treat patients with IBD.

## Introduction

Inflammatory bowel disease (IBD), mainly composed of Crohn’s disease (CD) and ulcerative colitis (UC), is a chronic, relapsing inflammatory disorder of the gastrointestinal tract, characterized by diarrhea, abdominal pain, weight loss, and increased risk of colorectal cancer ^1^. The rising prevalence of IBD in North American and around the world ^2^, coupled with the significant lifetime morbidity and financial burden ^3^, clearly highlights the urgent need for IBD research in order to identify novel therapeutic targets and develop potent and cost-effective treatments.

The intestinal epithelial cells (IECs) constitute a critical line of defense that plays a pivotal role in regulating host-microbiota interaction and intestinal homeostasis ^4^. Although IBD is a multifactorial disease, accumulating studies have linked mitochondrial dysfunction and oxidative stress within intestinal epithelia to IBD pathogenesis, suggesting a bioenergetics failure of the intestinal mitochondria in IBD, which consequently leads to excessive oxidative stress and tissue damage and results in disruption of intestinal homeostasis ^5, 6^. Mitochondrial biogenesis, the process by which new mitochondria are generated and repaired, plays a significant role in maintaining cellular metabolic homeostasis ^7^. Peroxisome proliferator-activated receptor-γ coactivator 1-α (PGC-1α) is the master transcriptional coactivator of genes encoding proteins responsible for the regulation of mitochondrial biogenesis and function ^8^. Recently, it has been reported that PGC-1α is highly expressed in intestinal epithelia ^9^, but significantly reduced in colitic mice and IBD patients ^10, 11^. Further, PGC-1α induction in IECs was shown to maintain mitochondrial integrity and barrier function and decrease intestinal inflammation ^10^.

The SET and MYND domain containing proteins (SMYDs), currently composed of five members (SMYD1-5), are a special class of protein methyltransferases that mediate lysine methylation of histones and non-histone proteins and are involved in transcriptional regulation and cellular signaling and function ^12, 13^. Among the five SMYD proteins, SMYD4 and SMYD5 are much less studied. SMYD5 has been reported to methylate histone H4 and regulate macrophage inflammation ^14^, embryonic stem cell self-renewal and differentiation ^15^, and zebrafish hematopoiesis and embryogenesis ^16^. So far, the only substrate that has been reported to be methylated by SMYD5 is histone H4 at lysine 20 ^14, 15^. However, the functional role of SMYD5 (especially intestine-specific) and the underlying mechanisms in regulating mitochondrial function and intestinal homeostasis in the context of health and disease (such as IBD) are completely unknown.

In this study, we have uncovered a novel non-histone target of SMYD5, PGC-1α, a critical transcriptional co-activator for mitochondrial biogenesis and function ^8^. Our study demonstrates that SMYD5 modulates mitochondrial functions in IECs through regulating the stability of PGC-1α in enzymatical activity dependent manner. Moreover, deficiency of Smyd5 in IECs protects mice from experimental colitis by preserving PGC-1α protein and promoting mitochondrial biogenesis and functions in IECs. Therefore, targeting SMYD5 expression in IECs may be a potential therapeutic strategy for IBD treatment.

## Materials and Methods

More detailed information is provided in the online **Supplemental Materials and Methods** section.

### Human Colonic Samples

Paraffin-embedded specimens of human colonic mucosa samples of control individuals (n = 8) and active Crohn’s disease patients (n = 8) were obtained from Biomax Company (Rockville, MD).

### Generation and Genotyping of IEC-specific Smyd5 Conditional KO Mice

To generate IEC-specific Smyd5 knockout mice (denoted Smyd5^ΔIEC^), embryonic stem (ES) cell clones with conditional potential targeting exon were obtained from the European Mouse Mutant Archive (EM:06942). Male ES cells were injected into C57BL/6 blastocysts at Georgia State University Transgenic and Gene Targeting Core facility. The targeted Smyd5 allele contains a LacZ reporter, the FLP-FRT sites, the neomycin-resistant marker, and the Cre-loxP sites flanking the exon 2 of murine Smyd5 gene (**Supplementary Figure 1**). The pups with the transmission of heterozygous Smyd5^fl/+^ was confirmed by PCR analysis. The mice carrying Smyd5^fl/+^ allele were mated till the Smyd5^fl/fl^ colony was obtained. Smyd5^fl/fl^ mice were then bred with mice expressing intestinal specific Cre-recombinase under the control of the villin promoter (catalog number 4586; The Jackson Laboratory) to generate both littermate control (Smyd5^fl/fl^) mice and IEC-specific Smyd5 KO (Smyd5^ΔIEC^) mice. Both Smyd5^fl/fl^ and Smyd5^ΔIEC^ mice were genotyped by PCR using primers listed in **Supplementary Table 1**. In this study, all the mice used were on C57BL/6 background, and the mutant lines were backcrossed for at least 6 generations and littermates were used as controls in all experiments. The animal studies were approved by the institutional animal care and use committee of Georgia State University.

### DSS-induced Colitis in Mice

Mice at 8-10 weeks of age were treated with 2.5% (w/v) dextran sodium sulfate (DSS, colitis grade, MW 36-50 kDa; MP Biochemicals) in drinking water *ad libitum* for 7 days and DSS-free water for another 2 days. In water control groups, mice were administered with DSS-free drinking water for 9 days. The severity of colitis was recorded daily and scored based on body weight loss, stool consistency, and presence of blood in the stools. On day 9, the colonic tissues were collected and fixed in 10% formaldehyde for 24 hrs, embedded in paraffin, sectioned for hematoxylin and eosin (H&E) or immunohistochemistry (IHC) staining. The disease activity index (DAI) and histological grade of colonic inflammation were evaluated as we previously reported ^17, 18^. The evaluation was performed by an experienced investigator who was blinded to the experimental design.

### Isolation of IECs

IECs were isolated as reported ^19^ with minor modifications. Briefly, colonic tissues were removed from the mice and opened longitudinally and cut into pieces followed by washing with PBS. IECs were then isolated following incubation of tissue pieces in PBS containing 30 mM EDTA and 2 mM DTT for 20 min at 37°C with gentle shaking at 200 rpm. Cells were then subjected to vigorous shaking for 30 s. After removal of the tissue debris, the isolated cells were pelleted by centrifugation at 1,000 g for 5 min.

### Mitochondria Isolation

Mitochondria were freshly isolated from IECs of Smyd5^fl/fl^ and Smyd5^ΔIEC^ mice as described before ^20^ using a mitochondria isolation kit (MITOISO1, Sigma) according to the manufacturer’s instruction.

### Oxygen Consumption Rate (OCR) Analysis

XFe96 extracellular flux analyzer (Seahorse Bioscience) was used to measure oxygen consumption rate (OCR) as previously described ^21^. Briefly, parental HCT116 cells and HCT116 cells with SMYD5 overexpression (OE) or knockout (KO) were seeded in 96-well plate (in triplicates) (#101085-004, Seahorse Bioscience) one day before measurement. On the day of measurement, the cells were incubated at 37°C and the medium was replaced with 180 μl XF assay medium (#102365-100, Seahorse Bioscience) containing 10 mM glucose, 1 mM pyruvate, and 2 mM glutamine at pH 7.4. Oligomycin, trifluoromethoxy carbonyl cyanide phenylhydrazone (FCCP), and rotenone/antimycin A provided with the XF Cell Mito Stress Test Kit (#103015-100, Seahorse Bioscience) were prepared in XF assay medium (100840-000, Seahorse Bioscience). Measurement was performed at 37°C to detect basal respiration, maximal respiration, proton leak, and coupled respiration. Data were analyzed using Wave software provided by Seahorse Bioscience.

### Intestinal Permeability Assay

Intestinal permeability assay was performed by evaluation of fluorescein isothiocyanate (FITC)-dextran (Sigma Aldrich) in the blood as previously described ^22^. Briefly, Smyd5^fl/fl^ and Smyd5^ΔIEC^ mice were subjected to DSS-induced acute colitis as described above. On the last day of DSS treatment, mice were fasted for 4 hrs and administered by oral gavage with FITC-dextran tracer (4 kDa, 0.4 mg/g body weight) dissolved in PBS. Hemolysis-free serum was then collected after 1 h and 4 hrs post-gavage. A standard curve for FITC-dextran concentration was established by serial dilution of known amount of FITC-dextran, and serum from mice administered with PBS only was used to determine the background signal. The samples were diluted in PBS and the absorbance at 520 nm was measured.

### Statistical Analysis

Data were expressed as mean ± standard error of the mean (SEM). Graphs representing means ± SEM were obtained from at least 3 independent experiments. Statistical analyses were performed in GraphPad Prism 8.0 (GraphPad Software, Inc., San Diego, CA). Pairwise comparisons were made using the unpaired Student’s *t* test, while 1-or 2-way ANOVA with Bonferroni’s post-hoc test was used for multiple comparisons, and asterisk and pound symbols were used to denote statistical significance (^*/#^*p* < 0.05; ^**/##^*p* < 0.01; ^***/###^*p* < 0.001). Pearson’s correlation coefficient was used to assess the relationship between the expression levels of SMYD5 and PGC-1α in colon samples from IBD patients and healthy controls.

## Results

### Clinical Relevance of SMYD5 and PGC-1α in Colon Samples from IBD Patients

Recent studies have shown that SMYD5 regulates expression of toll-like receptor 4 (TLR4) target genes including CXCL10, IL-1α, and CCL4 during immune response ^14^. Upon challenge with pro-inflammatory stimuli, such as lipopolysaccharide, the expression of SMYD5 was significantly up-regulated in macrophages ^14^, suggesting a critical role of SMYD5 in immunity and inflammatory diseases.

To investigate if SMYD5 is involved in IBD, which is characterized by chronic inflammation of the gastrointestinal tract, we first sought to examine its expression in human colonic tissues. Immunohistochemical (IHC) staining was performed in the colon sections from healthy subjects and IBD patients. The results showed that SMYD5 was mainly expressed in colonic epithelia (i.e., IECs) (**Figure 1A**). Notably, the expression level of SMYD5 was upregulated in colonic epithelia from IBD patients compared to that from healthy controls (**Figure 1A and 1B**, left) suggesting the involvement of SMYD5 in the pathological process of IBD. It is well established that mitochondrial dysfunction is associated with IBD ^23^, therefore we also examined the expression of PGC-1α, a critical transcription coactivator that regulates mitochondrial biogenesis and functions ^8^. Significant reduction of PGC-1α expression in IECs was observed in the colon sections from IBD patients versus healthy subjects (**Figure 1A and 1B**, right), which is consistent with previous study ^10^. In order to reveal the potential relationship between SMYD5 and PGC-1α expression levels, we plotted SMYD5 expression level against PGC-1α level in colonic epithelia from both healthy controls and IBD patients. Quantification of SMYD5 and PGC-1α expression in colonic epithelia revealed a significant inverse correlation between these two proteins (**Figure 1C**). These results clearly indicate that SMYD5 is upregulated in inflamed intestinal epithelia and may be involved in IBD pathogenesis.

**Figure 1.**
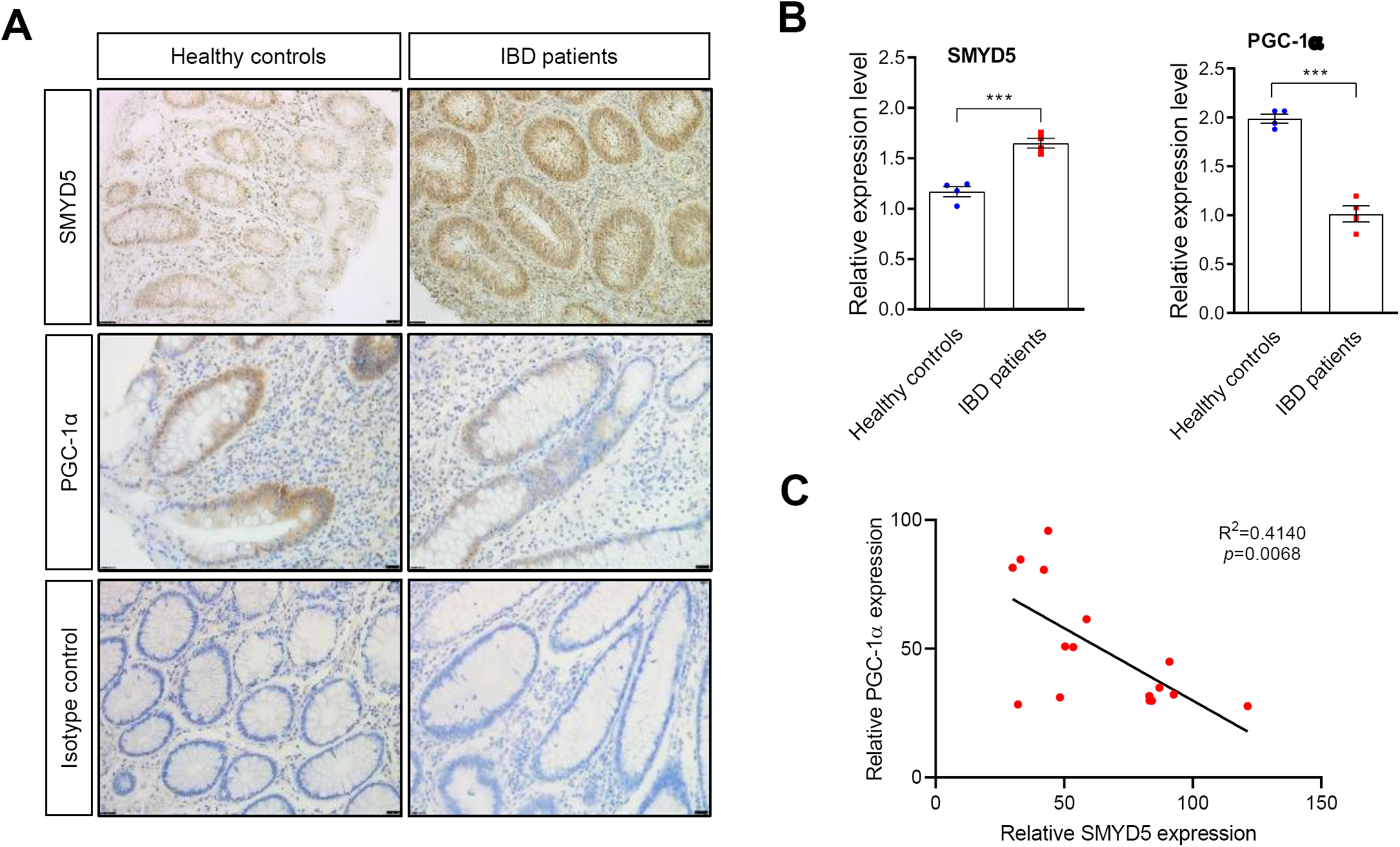
Clinical relevance of SMYD5 and PGC-1α in colonic samples from IBD patients. (A) Immunohistochemistry (IHC) staining of the expression of SMYD5 and PGC-1α in colon sections from healthy controls and IBD patients. Isotope controls with no primary antibodies are also shown (bottom panel). Scale bars, 50 μm. (B) Quantification of the IHC staining of SMYD5 and PGC-1α expression in IECs in colon sections from healthy controls and IBD patients. ^*\*\*\**^*p* < 0.001; n = 4. (C) Scatterplots between the relative expression (IHC staining intensity in arbitrary unit) of SMYD5 and PGC-1α in colonic epithelia from healthy controls and IBD patients. *p* = 0.0068; n = 16.

### SMYD5 Depletion in IECs Protects Mice from DSS-induced Experimental Colitis

To determine the potential importance of epithelial SMYD5 in IBD pathogenesis, we generated IEC-specific Smyd5 knockout mice. Smyd5 floxed mice (Smyd5^fl/fl^) were bred with Villin-Cre transgenic mice to produce IEC-specific Smyd5 conditional KO mice (Smyd5^ΔIEC^) (**Supplementary Figure 1**). IEC-specific ablation of Smyd5 gene was confirmed by PCR analysis (**Figure 2A**). Lack of SMYD5 protein expression specifically in IECs and intestinal epithelia was also validated by immunoblot analysis (**Figure 2B**, left) and immunofluorescence staining (**Figure 2B**, right), respectively.

**Figure 2.**
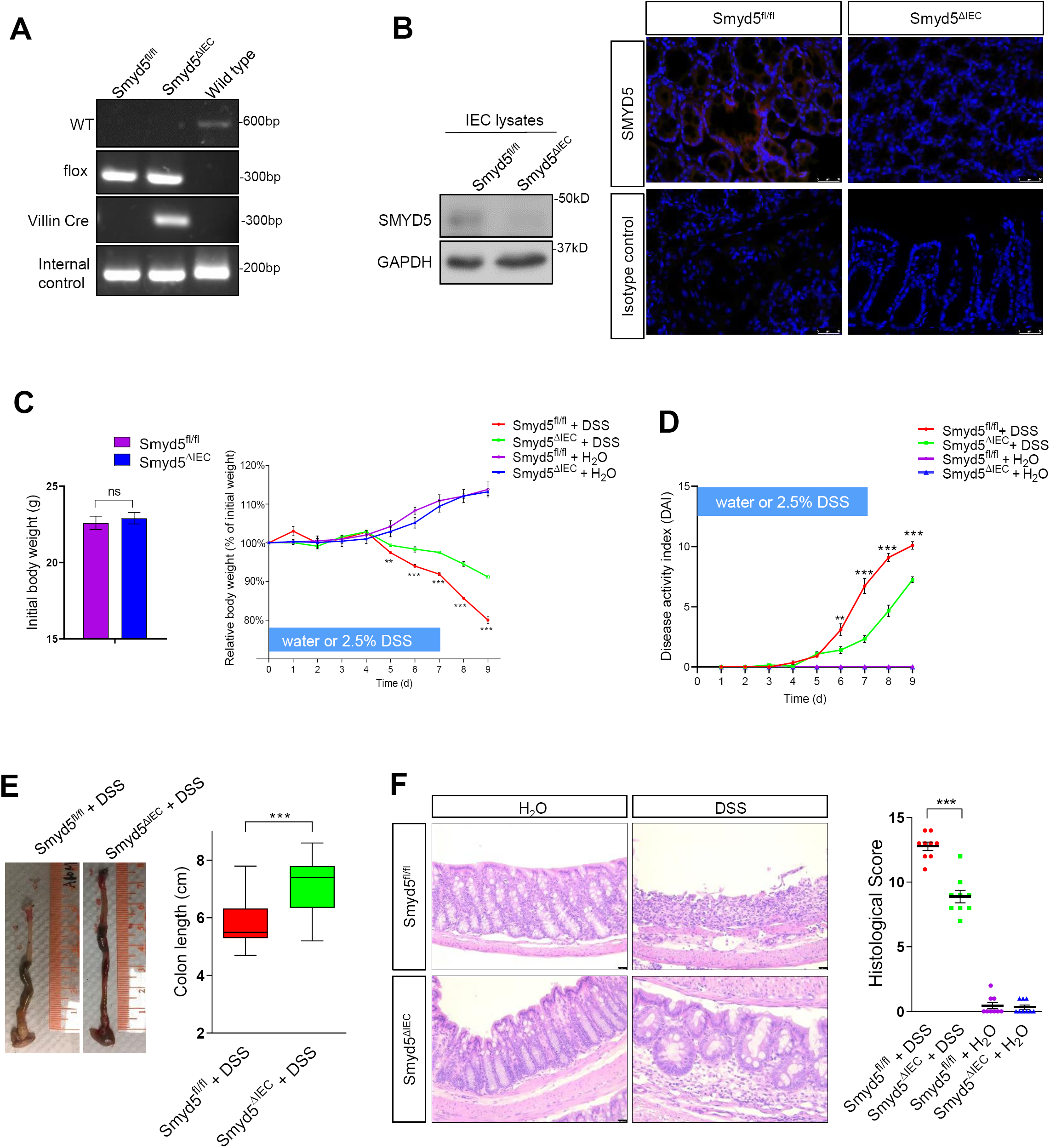
SMYD5 depletion in IECs protects mice from DSS-induced experimental colitis. (A) Genotyping of IEC-specific Smyd5 conditional KO mice (Smyd5^ΔIEC^) and Smyd5 floxed mice (Smyd5^fl/fl^) by PCR analysis. (B) Representative images of SMYD5 expression in murine IECs by immunoblot analysis (left panel) and in intestinal epithelia by immunofluorescence staining (right panel). Isotope controls with no primary antibody are also shown (right, bottom panel). Scale bars, 25 μm. (C) Initial body weight (left panel) and percentage body weight (relative to initial body weight) after DSS or water treatment (right panel) of Smyd5^fl/fl^ and Smyd5^ΔIEC^ mice. ^**^*p <* 0.01, ^*\*\*\**^*p* < 0.001, DSS-treated Smyd5^fl/fl^ versus DSS-treated Smyd5^ΔIEC^ mice, n = 12. (D) Changes in disease activity index (DAI). ^**^*p <* 0.01 on day 6, and ^***^ *p*< 0.001 on days 7 to 9, Smyd5^ΔIEC^ versus Smyd5^fl/fl^ mice upon DSS treatment, n = 12. (E) Gross morphology of representative colons (left) and colon length measurements (right) in Smyd5^fl/fl^ and Smyd5^ΔIEC^ mice after DSS administration. ^*\*\*\**^*p* < 0.001, n = 20. (F) Representative H&E staining (left) and histology scores (right) for colon sections from Smyd5^fl/fl^ and Smyd5^ΔIEC^ mice after DSS or water treatment. Scale bars, 50 μm. ^*\*\*\**^*p* < 0.001, n = 9.

To investigate whether Smyd5 deficiency affects the development of IBD, both floxed and KO mice were subjected to drinking water supplemented with 2.5% DSS, a colitogenic chemical widely used to induce colitis in experimental animals ^24^. In control groups, all mice were provided with regular drinking water for 9 days. In DSS groups, mice were administered with DSS containing water for 7 consecutive days and then regular drinking water for additional 2 days. Smyd5^fl/fl^ and Smyd5^ΔIEC^ mice administered with water only had gained weight to a similar extent (**Figure 2C**). DSS administration damaged the colonic mucosal barrier, leading to intestinal inflammation and weight loss in both Smyd5^fl/fl^ and Smyd5^ΔIEC^ mice. Although both mouse lines lost weight after DSS administration, Smyd5^ΔIEC^ mice had significantly less weight loss than Smyd5^fl/fl^ mice during days 5 to 9, with an estimated weight change difference of 11% at day 9 between the two mouse lines (**Figure 2C**). Furthermore, Smyd5^ΔIEC^ mice had better-shaped (less loose) stools and less blood in their stools than Smyd5^fl/fl^ mice during days 6 to 9. These differences resulted in a lower disease activity index (DAI) (**Figure 2D**). Consistent with lower DAI scores, Smyd5^ΔIEC^ mice exhibited less colon shortening upon exposure to DSS, as compared with Smyd5^fl/fl^ mice (**Figure 2E**). Histological analysis on day 9 post DSS administration revealed less extensive ulceration and erosion, and less severe inflammatory cell infiltration, as well as less thickening of the mucosa with edema in the colon sections of Smyd5^ΔIEC^ mice compared with that of Smyd5^fl/fl^ mice, leading to lower histological scores (**Figure 2F**). Together, these data demonstrate that IEC-specific Smyd5 deficiency reduces the severity of DSS-induced colitis in mice.

### SMYD5 Negatively Regulates PGC-1α Expression at Post-transcriptional Level

Accumulating studies have demonstrated that members of SMYD family not only mediate the methylation of histone proteins but also non-histone substrates ^13^. Moreover, SMYD family of methyltransferases have been shown to modulate protein expression at both transcriptional and post-transcriptional levels ^12, 13, 25^. Based on the above observations, it is possible that SMYD5 may also be involved in the regulation of PGC-1α expression in IECs and subsequently the modulation of mitochondrial function. In order to further reveal the relationship between SMYD5 and PGC-1α, we altered the expression of SMYD5 in a human colonic epithelial cell line HCT116 by overexpressing or depleting SMYD5 and examined PGC-1α expression via immunofluorescence staining and immunoblotting analysis. The results showed that PGC-1α was decreased when SMYD5 was overexpressed; conversely, SMYD5 KO resulted in up-regulation of PGC-1α (**Figure 3A and B**). This indicates that SMYD5 negatively regulates the expression of PGC-1α in IECs.

**Figure 3.**
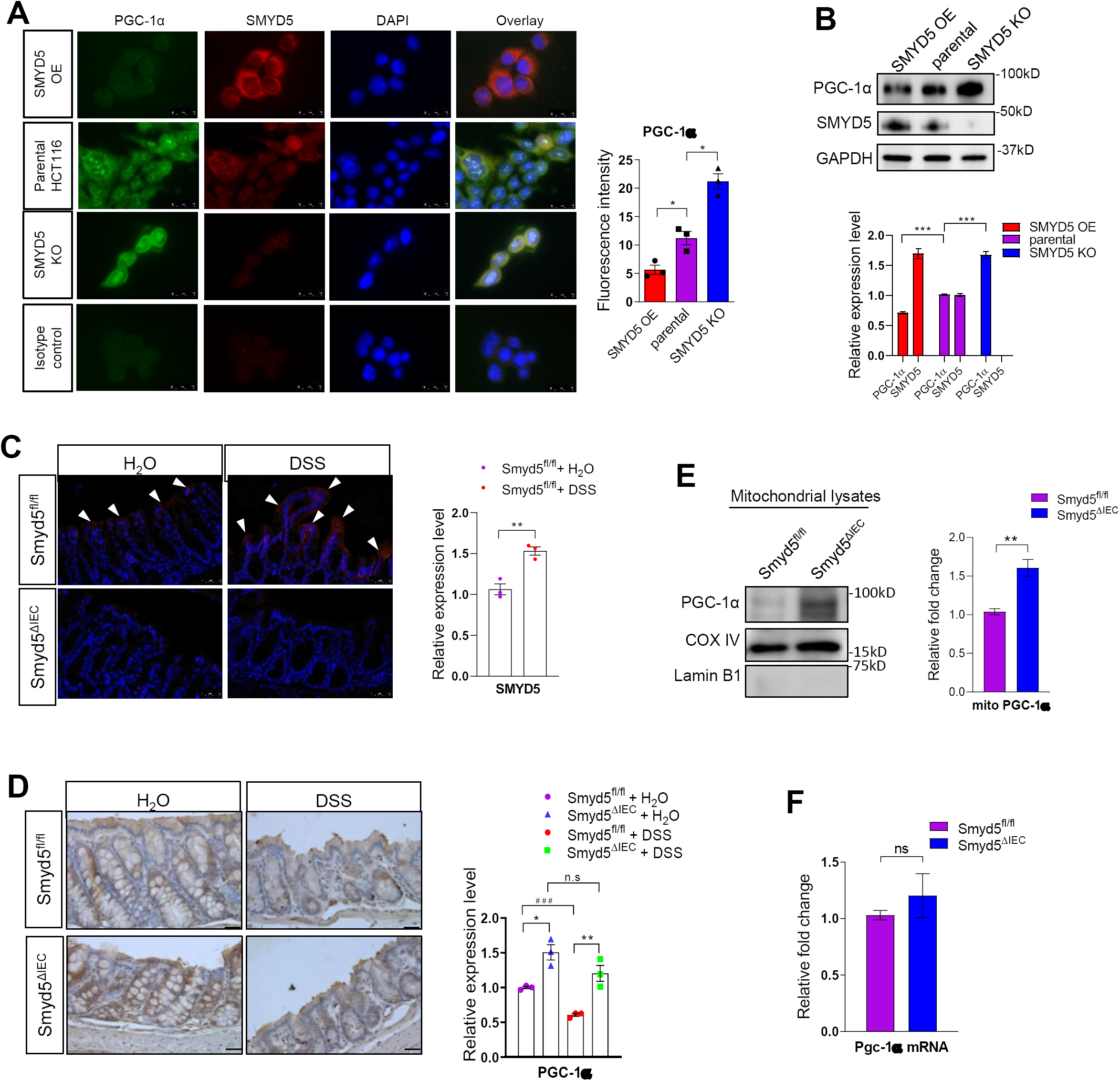
SMYD5 negatively regulates PGC-1α at post-transcriptional level. (A) Representative images (left) and quantitative analysis (right) of immunofluorescence (IF) staining of SMYD5 and PGC-1α expression in HCT116 cells with either SMYD5 overexpressed (OE) or knocked out (KO). Isotope controls with no primary antibodies are also shown (left, bottom panel). Scale bars, 25 μm. ^*\**^*p* < 0.05, n = 3. (B) Representative Western blots (top) and quantitative analysis (bottom) of SMYD5 and PGC-1α expression (normalized to GAPDH) in HCT116 cells with either SMYD5 OE or SMYD5 KO. The expression levels of SMYD5 and PGC-1α in parental HCT116 cells were set as 1. ^***^*p* < 0.001, comparison of PGC-1α levels among groups; n = 3. (C) Representative IF staining (left) and quantitative analysis (right) of SMYD5 (red color indicated by white arrowheads) in epithelia of the intestinal mucosa in Smyd5^fl/fl^ mice following water or DSS administration. Note: SMYD5 was not detected in Smyd5^ΔIEC^ mice. Scale bars, 50 μm. ^*\*\**^*p* < 0.01, n = 3. (D) Representative IHC staining (left) and quantitative analysis (right) of PGC-1α (brown color) in intestinal epithelia from Smyd5^fl/fl^ and Smyd5^ΔIEC^ mice following water or DSS administration. Scale bars, 50 μm. DSS vs. water-treated ^###^*p* < 0.001, Smyd5^ΔIEC^ vs. Smyd5^fl/fl *^*p* < 0.05, ^**^*p* < 0.01, n.s., not significant; n = 3. (E) The mitochondria were isolated from IECs of Smyd5^fl/fl^ and Smyd5^ΔIEC^ mice followed by detection of mitochondrial PGC-1α protein level via immunoblot analysis. COX IV and Lamin B1 were used as specific markers for mitochondrial and nuclear fractions, respectively. Mitochondrial PGC-1α expression was quantified and presented as the fold change relative to the level in Smyd5^fl/fl^ IECs. ^**^*p* < 0.01, n = 3. (F)Transcription level (mRNA) of Pgc-1α was assessed by RT-qPCR (presented as relative fold change) in IECs isolated from Smyd5^fl/fl^ and Smyd5^ΔIEC^ mice. *p* = 0.4346, n = 3; n.s, not significant.

Interestingly, similar changes of SMYD5 and PGC-1α expression were also observed in the murine model of colitis demonstrated by immunofluorescence staining (**Figure 3C**) and immunohistochemistry (**Figure 3D**). Immunofluorescence staining showed that DSS administration resulted in significantly up-regulated SMYD5 in epithelia of inflamed mucosa of Smyd5^fl/fl^ control mice compared to water administration (**Figure 3C**, top panel) as reflected by elevated fluorescence intensity (red color as indicated by white arrowheads). In parallel, the expression of PGC-1α in epithelial mucosa of the intestine was decreased in both Smyd5^fl/fl^ and Smyd5^ΔIEC^ mice with DSS-induced colitis compared to water treatment (**Figure 3D**), which is consistent with previous study ^10^. Of note, PGC-1α expression level in intestinal epithelia of Smyd5^ΔIEC^ mice was significantly higher than that from Smyd5^fl/fl^ mice at both baseline (water treatment) and upon DSS-colitis induction (**Figure 3D**). This is consistent with the results observed from human colon sections (**Figure 1A and B**). To further confirm these findings, HCT116 cells were treated with TNF-α or IFN-γ, the proinflammatory cytokines that have been reported to be up-regulated in intestinal mucosa in human IBD and murine experimental colitis ^26, 27^, and the expression of SMYD5 and PGC-1α was determined by immunoblot analysis. The results showed that treatment with the proinflammatory cytokines resulted in up-regulated SMYD5 and down-regulated PGC-1α (**Supplementary Figure 2**).

PGC-1α has been shown to reside in mitochondria and regulate mitochondrial DNA (mtDNA) biogenesis ^28^. Thus, we asked if SMYD5 is involved in regulating mitochondrially-located PGC-1α protein level. We isolated mitochondria from IECs of Smyd5^fl/fl^ and Smyd5^ΔIEC^ mice as described before ^20^ and examined PGC-1α protein level by immunoblot analysis. The results showed that the expression of mitochondrial PGC-1α in IECs from Smyd5^ΔIEC^ mice is significantly higher than that from Smyd5^fl/fl^ mice (**Figure 3E**). Further, RT-qPCR analysis showed that there is no significant difference in the transcriptional levels of PGC-1α in IECs from Smyd5^fl/fl^ versus Smyd5^ΔIEC^ mice (**Figure 3F**) suggesting that SMYD5 regulates PGC-1α protein level at the post-transcriptional level.

### SMYD5 Is Critically Involved in Regulating Mitochondrial Function and Intestinal Barrier Integrity

PGC-1α is a multifunctional transcriptional co-activator critically involved in mitochondrial biogenesis and oxidative phosphorylation to modulate diverse cellular functions and processes ^28, 29^. Accumulating evidence from recent studies has implicated mitochondrial dysfunction within the intestinal epithelium in the onset and progression of IBD ^5, 6, 10^. We speculated that up-regulation of SMYD5 in IECs during IBD may result in the decrease of PGC-1α protein level, leading to attenuated mitochondrial biogenesis and functions. To test this hypothesis, we examined the expression of markers of mitochondrial biogenesis via RT-qPCR using IECs from Smyd5^fl/fl^ and Smyd5^ΔIEC^ mice. The results showed that Smyd5 deficiency led to up-regulated mitochondrial biogenesis as revealed by the significantly elevated expression of mitochondrial transcription factor A (Tfam) that controls mtDNA replication and transcription and thus promotes mitochondrial biogenesis ^30^, and cytochrome c oxidase (COX) I which is a mtDNA-encoded protein ^31^. (**Figure 4A**). To further confirm this observation, we also examined the abundance of mtDNA content and the expression of mtDNA-encoded genes (COX I and COX II) in HCT116 cells with overexpressing or knockout SMYD5. RT-qPCR analyses revealed that SMYD5 knockout (KO) in HCT116 cells significantly increased mtDNA content as compared to control or parental cells (**Supplementary Figure 3A**). Moreover, SMYD5 overexpression (OE) decreased, while SMYD5 KO increased, the transcriptional levels of both COX I and COX II in HCT116 cells (**Supplementary Figure 3B and 3C**). We then determined the mitochondrial counts in IECs from Smyd5^fl/fl^ and Smyd5^ΔIEC^ mice using transmission electron microscopy (TEM) as described before ^29^. The data demonstrated that Smyd5 depletion resulted in a slight but significant elevation in number of mitochondria in IECs (**Figure 4B**).

**Figure 4.**
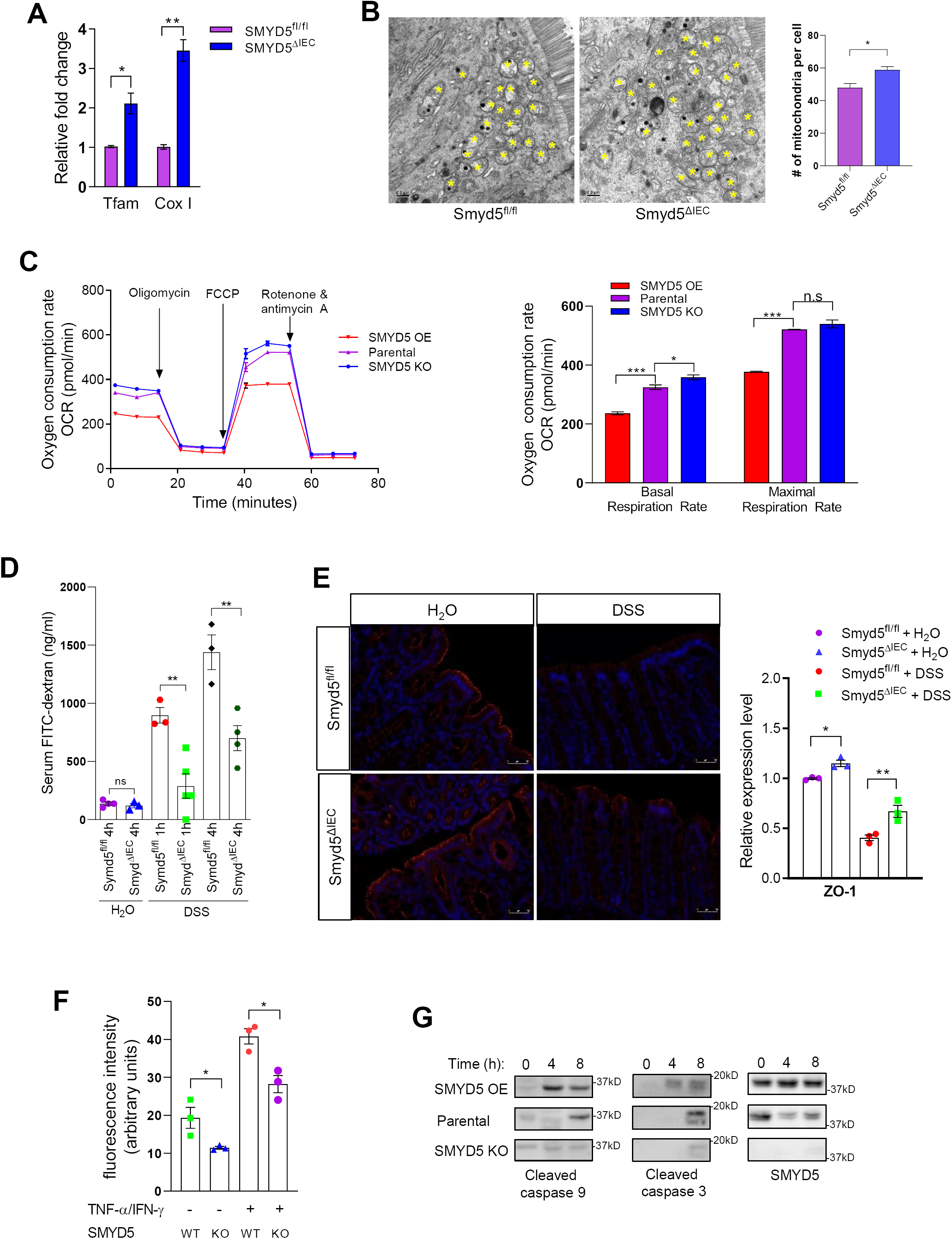
Smyd5 is critically involved in regulating mitochondrial function and intestinal barrier integrity. (A) RT-qPCR analysis of mRNA (presented as relative fold change) of mitochondrial biogenesis markers Tfam (^*\**^*p* < 0.05, n = 3) and Cox I (^*\*\**^*p* < 0.01, n = 3) in IECs isolated from Smyd5^fl/fl^ and Smyd5^ΔIEC^ mice. (B) Representative TEM images (left) of IECs from Smyd5^fl/fl^ and Smyd5^ΔIEC^ mice and quantitative analyses of number of mitochondria per cell (right; ^*\**^*p* < 0.05, n = 3). Yellow asterisks (*) indicate mitochondria. Scale bars: 500 nm. (C) Oxygen consumption rates (OCR) measured by Seahorse XF96 analyzer (left) in HCT116 cells with SMYD5 overexpression, unchanged (parental), or knockout. Glucose (25 mmol/L) and pyruvate (1 mmol/L) were supplied as substrates. Quantitation of basal and maximal respiration capacity (right). ^*^*p* < 0.05, ^***^*p* < 0.001, n.s., not significant; n = 3 (D) Smyd5^fl/fl^ and Smyd5^ΔIEC^ mice were subjected to DSS-induced acute colitis for 7 days. On the last day of DSS feeding, mice were fasted overnight and gavaged with FITC-dextran (4 kDa) and blood was collected for measurement of FITC-dextran at 1 h or 4 hrs post-gavage. Intestinal permeability was measured by the concentration of FITC-dextran in the blood serum. ^\**\**^*p* < 0.01, n = 3-5 per group. (E) Representative IF staining of ZO-1 expression (left) in colon tissues of Smyd5^ΔIEC^ or Smyd5^fl/fl^ mice administered with water or DSS. Red fluorescence: ZO-1; blue fluorescence: DAPI. Scale bars: 50 μm. Quantification of ZO-1 staining (right) in colon tissues of water or DSS-administered Smyd5^fl/fl^ or Smyd5^ΔIEC^ mice. ^*^*p* < 0.05, ^\**\**^*p* < 0.01; n = 3. (F) The cellular oxidative stress was measured using a fluorescence-based assay as described in Methods without or with exposure to proinflammatory cytokines (20 ng/ml TNF-α and 20 ng/ml IFN-γ, 24 hrs) in SMYD5 KO HCT116 cells and parental HCT116 cells (SMYD5 WT). ^*^*p* < 0.05, n = 3. (G) Parental HCT116, SMYD5 OE HCT116, or SMYD5 KO HCT116 cells were treated with H_2_O_2_ (100 µM) for 4 or 8 hrs and the expression of cleaved caspase 9 and 3 as well as SMYD5 was examined by Western blot analysis using respective antibodies.

It has been reported that augmented oxidative phosphorylation and increased intestinal ATP protected mice from DSS and trinitrobenzene sulfonate (TNBS)-induced experimental colitis ^32^. We next examined the effect of SMYD5 expression on mitochondrial respiratory function of IECs. HCT116 cells with SMYD5 KO or OE were used for Seahorse assays to measure oxygen consumption rate (OCR), which reflects mitochondrial function ^20, 33^. Oxygen consumption was measured under basal condition, following the sequential addition of oligomycin, the pharmacological uncoupler FCCP, the Complex III and I inhibitors antimycin A and rotenone (**Figure 4C**, left). SMYD5 OE significantly reduced the basal OCR and the maximal respiratory capacity after FCCP treatment, while SMYD5 KO led to the opposite effect (**Figure 4C**, right).

One of the characteristic features of IBD is loss of intestinal mucosal barrier integrity ^34^. Epithelial mitochondrial dysfunction has been implicated as a predisposing factor for compromised barrier integrity and increased gut permeability, perpetuating intestinal inflammation ^23^. Thus, it is possible that SMYD5 is involved in modulating epithelial monolayer integrity by regulating mitochondrial function of IECs. Toward this, we performed an *in vivo* intestinal permeability assay in water or DSS-administered Smyd5^fl/fl^ and Smyd5^ΔIEC^ mice, by oral gavage of FITC-dextran and measurement of FITC-dextran in the serum after 1 h or 4 hrs post-gavage. The results demonstrated that at basal condition without DSS injury (water administration), Smyd5^fl/fl^ and Smyd5^ΔIEC^ mice maintained a comparable barrier integrity reflected by low levels of plasma FITC-dextran in both strains (**Figure 4D**). DSS administration disrupted intestinal barrier integrity in both mouse strains as revealed by the drastic increase of serum FITC-dextran (namely, leaking of FITC-dextran into blood) in a time-dependent manner. However, serum FITC-dextran in Smyd5^ΔIEC^ mice was significantly lower than that in Smyd5^fl/fl^ mice suggesting a protective role of Smyd5 ablation in epithelial barrier integrity (**Figure 4D**). To investigate the mechanism by which SMYD5 deficiency might alter epithelial permeability in colitic mice, we examined the expression of tight junction proteins involved in intestinal epithelial barrier and transport functions. Immunofluorescence staining of ZO-1 protein in intestinal epithelia demonstrated a slight increase at baseline (water) in Smyd5^ΔIEC^ mice versus Smyd5^fl/fl^ mice (**Figure 4E**). Upon DSS challenge, ZO-1 staining was drastically decreased in intestinal epithelia of both Smyd5^fl/fl^ and Smyd5^ΔIEC^ mice, but was less decreased in Smyd5^ΔIEC^ as compared to Smyd5^fl/fl^ mice (**Figure 4E**).

Cellular oxidative stress induced by proinflammatory cytokines was evaluated in SMYD5 KO HCT116 cells and parental cells using a fluorescence-based assay ^35^. The results demonstrated that SMYD5 deficiency reduced the oxidative stress in cells at basal level, and exposure to proinflammatory cytokines (TNF-α and IFN-γ) increased cellular oxidative stress in both groups as revealed by increased fluorescence intensity (**Figure 4F**). However, SMYD5 depletion attenuated the elevation of cellular oxidative stress induced by proinflammatory cytokines (**Figure 4F**). Immunoblot analysis also demonstrated that SMYD5 overexpression in HCT116 cells exaggerated apoptosis (indicated by time-dependent increase in cleaved caspase 3 and 9 levels) as compared to HCT116 parental cells, while SMDY5 deletion reduced apoptotic response in HCT116 cells (**Figure 4G**). Collectively, these results clearly indicate that SMYD5 modulates intestinal barrier integrity by regulating mitochondrial functions, such as mitochondrial biogenesis and respiration, mitochondria-mediated apoptosis, probably through modulating epithelial PGC-1α protein level.

### PGC-1α Is a Substrate of SMYD5 Methyltransferase

Accumulating studies suggest that protein lysine methylation has been linked to their proteasomal degradation ^36-38^. For instance, methylation of lysine (K) 185 of E2F1 induced its proteasomal degradation, thus inhibited E2F1 apoptotic activity ^36^. In order to determine whether SMYD5 mediates methylation of PGC-1α to regulate its proteasomal degradation, we first evaluated the potential physical interaction between SMYD5 and PGC-1α. HEK293T cells were co-transfected with hemagglutinin (HA)-tagged SMYD5 (HA-SMYD5) and Flag-tagged PGC-1α (Flag-PGC-1α) and then subjected to co-immunoprecipitation assay. The results demonstrated that PGC-1α was co-immunoprecipitated with SMYD5 (**Figure 5A**).

**Figure 5.**
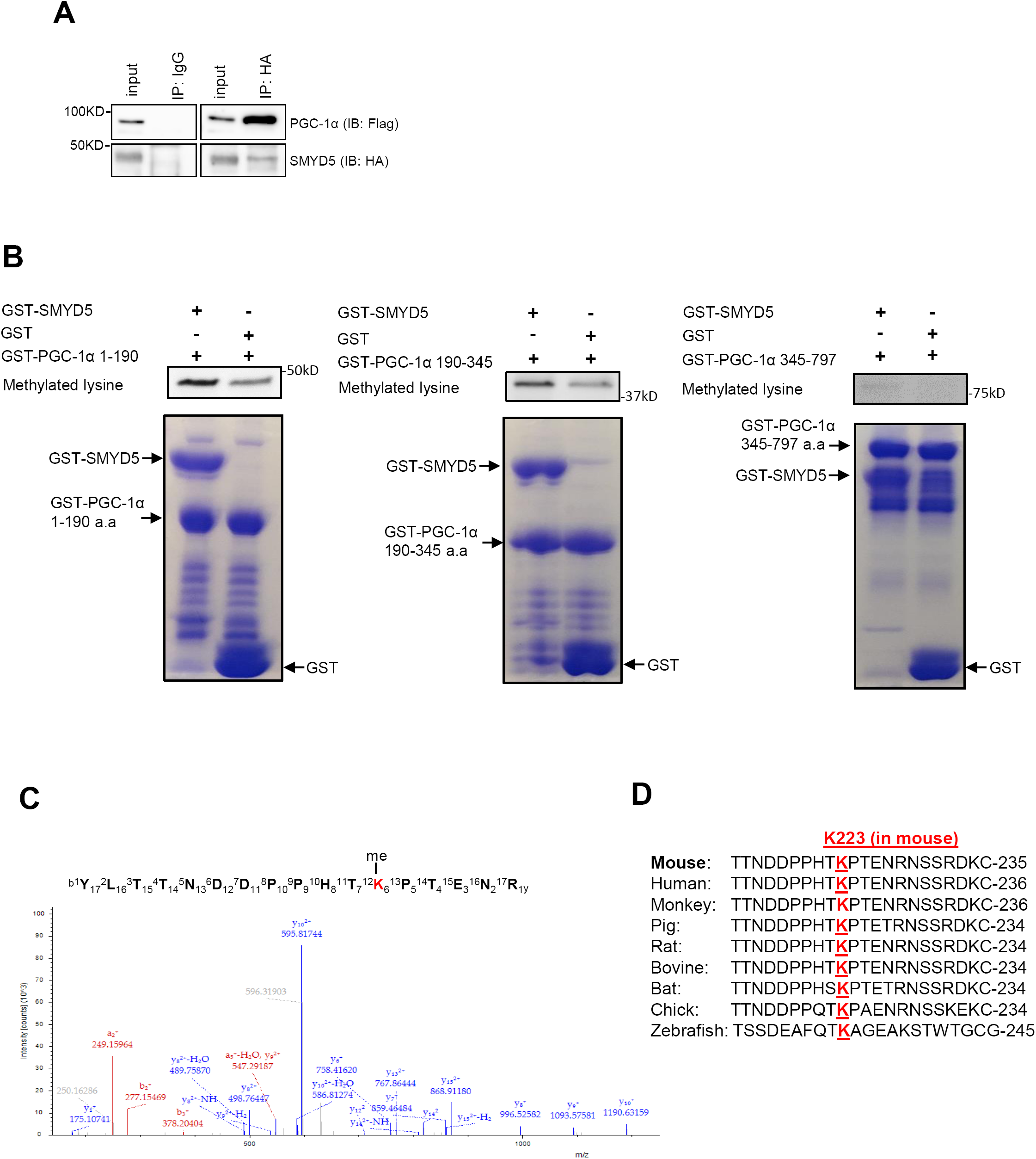
PGC-1α can be methylated by SMYD5. (A) Co-immunoprecipitation and immunoblot analysis of SMYD5-PGC-1α interaction in HEK293T cells co-transfected with HA-SMYD5 and Flag-PGC-1α. Whole cell lysates were immunoprecipitated with anti-HA or control IgG, and the immunocomplexes were immunoblotted using anti-Flag (for PGC-1α) and anti-HA (for SMYD5) antibodies. (B) *In vitro* methyltransferase assay of lysine methylation of PGC-1α and immunoblot analysis using a methyl lysine-specific antibody to detect methylated lysine(s) of PGC-1α. Bacterially purified GST-SMYD5 or GST alone was mixed with GST-PGC-1α fragments (GST-PGC-1α aa 1-190, GST-PGC-1α aa 190-345, and GST-PGC-1α aa 345-797) in a methylation reaction as described in the Methods. Bottom: the Coomassie blue staining showing equal amounts of GST-PGC-1α fragments mixed with either GST-SMYD5 protein or GST alone used in the in vitro methylation assay. (C) LC-MS/MS spectrum of the mono-methylated lysine 223 (K223) of murine PGC-1α fragment/peptide by SMYD5. The *in vitro* methylation assay was conducted as described in (B), followed by SDS-PAGE. LC-MS/MS analysis was conducted after digestion of samples by trypsin. (D) Alignment of peptide sequences spanning PGC-1a K223 (murine) among the indicated species.

In order to investigate if SMYD5 can methylate PGC-1α *in vitro*, GST-tagged SMYD5 (GST-SMYD5) and GST-tagged murine PGC-1α fragments (GST-PGC-1α aa 1-190, GST-PGC-1α aa 190-345, and GST-PGC-1α aa 345-797) were purified. GST-SMYD5 was mixed with GST-PGC-1α fragments or GST protein in a luminescence-based *in vitro* methyltransferase assay, a nonradioactive and antibody-free assay to determine enzyme activity of diverse families of methyltransferases ^39, 40^. As compared to GST alone, all GST-PGC-1α fragments demonstrated significantly increased methylation (as reflected by elevated luminescence signal) in the presence of SMYD5 (**Supplementary Figure 4A**). Because SMYD5 is known as a lysine methyltransferase, we next investigated if SMYD5 mediates methylation of lysine residues in PGC-1α protein. Purified GST-PGC-1α fragments (PGC-1α aa 1-190, aa 190-345, and aa 345-797) were mixed with GST-SMYD5 or GST alone in an *in vitro* methylation assay as reported before ^41^ followed by immunoblotting analysis using a methyl lysine-specific antibody to detect methylated lysine(s) of PGC-1α. The results showed that, compared to GST alone, GST-SMYD5 led to enhanced lysine methylation in PGC-1α aa 1-190 (**Figure 5B**, left) and PGC-1α aa 190-345 (**Figure 5B**, middle). Of note, PGC-1α fragments also demonstrated basal lysine methylation signal even in the absence of GST-SMYD5 (**Figure 5B**).

To identify the specific lysine residues that have been methylated by SMYD5, the *in vitro* methylation assay was conducted as described above, and mass spectrometric analysis was performed to identify the methylated lysine residue in all the PGC-1α fragments. The results showed that SMYD5 resulted in mono-methylation of lysine 223 (K223) of murine PGC-1α (**Figure 5C**, and **Supplementary Figure 4B**), an evolutionally conserved lysine residue among different species (from zebrafish to human) (**Figure 5D**). We could not detect any lysine methylation within fragments of PGC-1α aa 1-190 or PGC-1α aa 345-797. Taken together, these results suggest that SMYD5 mediates lysine methylation of PGC-1α at K223 and may modulate its degradation.

### SMYD5 Mediates Proteasomal Degradation of PGC-1α

There are two major fundamentally distinct mechanisms by which proteins are degraded in eukaryotic cells: the ubiquitin-proteasome pathway and the autophagy-lysosomal pathway. Previous studies have shown that PGC-1α is a short-lived, unstable protein primarily targeted for ubiquitin-proteasome dependent degradation at basal level ^42-45^. We next sought to explore how PGC-1α protein level is regulated by SMYD5. In alignment with previous studies ^43^, treatment with proteasomal inhibitor MG132 resulted in significant up-regulation of PGC-1α protein level in HEK293T cells (**Figure 6A**). However, blockade of the autophagy-lysosomal degradation pathway with bafilomycin A1 (an inhibitor of vacuolar H^+^-ATPase) and chloroquine (which inhibits autophagic flux by decreasing autophagosome-lysosome fusion) ^46^, did not affect the expression level of PGC-1α protein in HCT116 cells (**Supplementary Figure 5**), suggesting that PGC-1α degradation is mediated by the ubiquitin-proteasome pathway.

**Figure 6.**
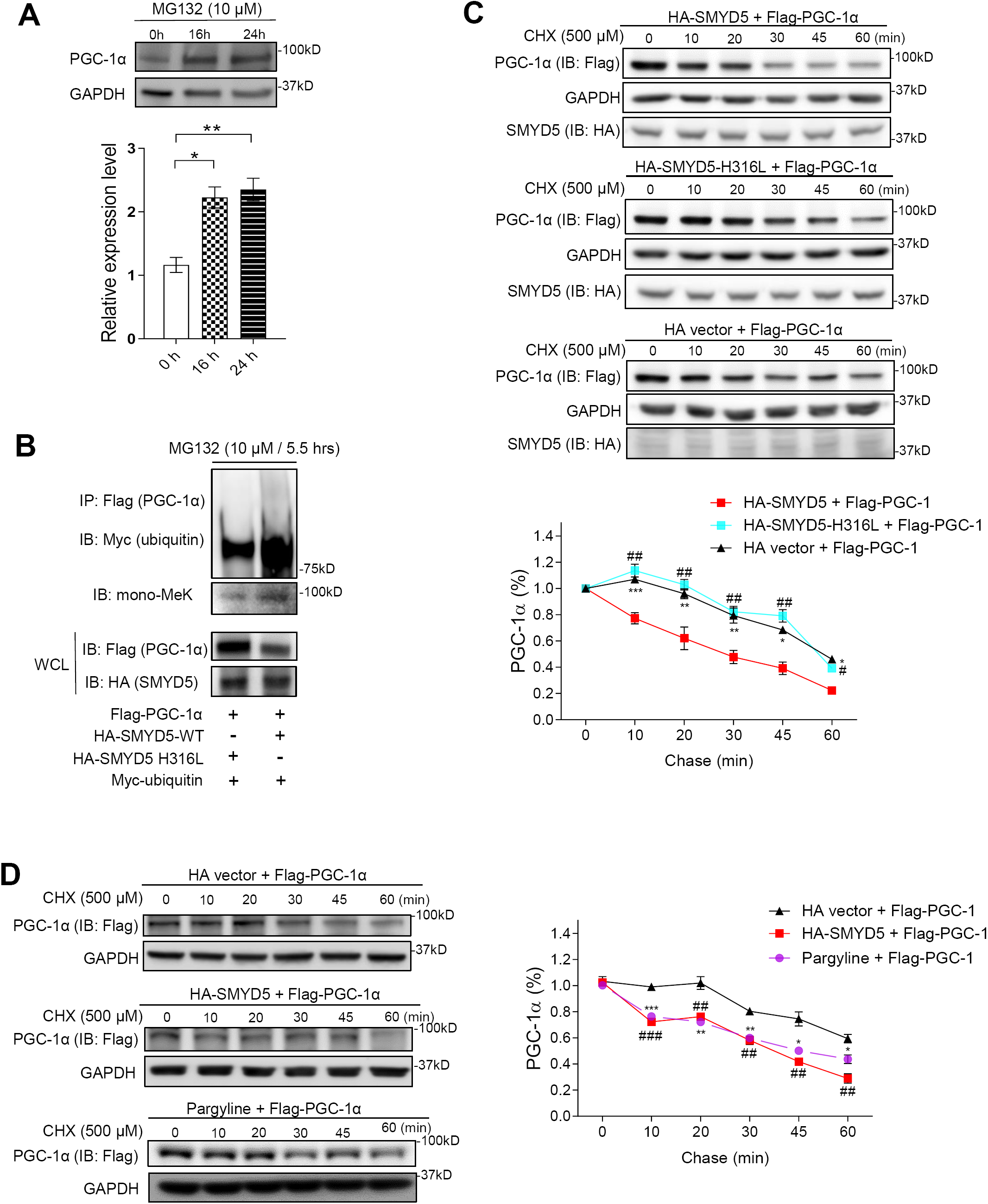
SMYD5 regulates proteasomal degradation of PGC-1α. (A) HEK293T cells were treated with the proteasome inhibitor MG132 (10 μM) for indicated times and total cell extracts were analyzed by Western blotting for PGC-1α protein amounts. ^*^*p* < 0.05, ^**^*p* < 0.01; n = 3. (B) HEK293T cells were co-transfected with myc-tagged ubiquitin (myc-ubiquitin), Flag-tagged PGC-1α (Flag-PGC-1α), and HA-tagged wildtype SMYD5 (HA-SMYD5-WT) or enzymatically inactive SMYD5 (HA-SMYD5-H316L) followed by treatment with MG132 (10 μM) for 5.5 hrs. Then whole cell lysates were subjected to immunoprecipitation with anti-Flag antibody (for Flag-PGC-1α) and the immunoprecipitated PGC-1α were analyzed by Western blot for ubiquitination using anti-myc antibody (for myc-ubiquitin) and for lysine methylation using anti-mono methyl lysine (mono-MeK) antibody. A small fraction of whole cell lysates (WCL) before immunoprecipitation was also immunoblotted for PGC-1α (anti-Flag) and SMYD5 (anti-HA). (C) HEK293T cells were transfected with Flag-PGC-1α along with HA-SMYD5-WT, or HA-SMYD5-H316L, or HA alone vector. Then cells were treated with cycloheximide (CHX, 500 μM) for indicated time periods before whole cell lysates were immunoblotted for PGC-1α (with anti-Flag antibody), SMYD5 (with anti-HA antibody), and GAPDH. The relative levels of Flag-PGC-1α protein at each time point (normalized to GAPDH) is plotted as a percentage (%) of the amount at 0 min. HA-SMYD5 vs. HA vector, ^*^*p* < 0.05, ^**^*p* < 0.01, ^***^*p* < 0.001; HA-SMYD5 vs. HA-SMYD5-H316L, ^#^*p* < 0.05, ^##^*p* < 0.01; n = 3. (D) HEK293T cells were transfected with Flag-PGC-1α along with HA alone vector or HA-SMYD5-WT. In some groups, cells transfected with Flag-PGC-1α were treated with lysine-specific demethylase 1 (LSD1) inhibitor, pargyline (2.5 mM) for 24 hrs. Then cells were treated with cycloheximide (500 μM) for indicated time periods before whole cell lysates were immunoblotted for PGC-1α (with anti-Flag antibody) and GAPDH. The relative levels of Flag-PGC-1α protein at each time point (normalized to GAPDH) is plotted as a percentage (%) of the amount at 0 min. HA vector vs. pargyline treatment, ^*^*p* < 0.05, ^**^*p* < 0.01, ^***^*p* < 0.001; HA vector vs. HA-SMYD5, ^##^*p* < 0.01, ^###^*p* < 0.001; n = 3.

Since proteasomal degradation is mediated by protein ubiquitination ^47^, we next investigated whether SMYD5-mediated methylation affects PGC-1α ubiquitination and degradation. HEK293T cells were co-expressed with myc-tagged ubiquitin (myc-ubiquitin), Flag-tagged PGC-1α (Flag-PGC-1α), and HA-tagged wildtype SMYD5 (HA-SMYD5-WT) or enzymatically inactive SMYD5 (HA-SMYD5-H316L) ^14^ and the ubiquitinated PGC-1α and total PGC-1α were examined by Western blot. The results showed that wildtype SMYD5 (HA-SMYD5-WT) significantly increased the lysine methylation and ubiquitination of PGC-1α compared with enzymatically inactive SMYD5 (HA-SMYD5-H316L) (**Figure 6B**). Of note, total PGC-1α amount (in whole cell lysates) was substantially reduced in cells co-expressed with HA-SMYD5-WT versus inactive mutant HA-SMYD5-H316L, even in the presence of proteasomal inhibitor MG132 (for 5.5 hrs) (lower band, **Figure 6B**). So far there have been a few E3 ligases, such as SCF^Cdc4^ (FBXW7) ^42, 45^ and RNF34 ^44^, that have been demonstrated to mediate PGC-1α ubiquitination and degradation. However, knocking down the expression of FBXW7 or RNF34 in IECs did not cause any significant change in PGC-1α protein levels (**Supplementary Figure 6**), suggesting that FBXW7 and RNF34 are not involved in the regulation of PGC-1α stability in IECs.

Next, we monitored the half-life of PGC-1α protein by pulse-chase analysis. HEK293T cells were transfected with Flag-PGC-1α, together with HA-SMYD5-WT, or HA-SMYD5-H316L, or HA alone vector, respectively, and the cells were treated with translation elongation inhibitor cycloheximide (CHX) to block protein synthesis, followed by chasing the remaining PGC-1α. Co-expression of wildtype SMYD5 with PGC-1α significantly reduced the half-life of PGC-1α protein (**Figure 6C**). However, co-expressing the enzymatically inactive SMYD5 mutant (HA-SMYD5-H316L) with PGC-1α demonstrated similar effect on PGC-1α half-life as HA alone vector (**Figure 6C**), indicating that the reduction of PGC-1α half-life by SMYD5 overexpression is SMYD5 enzymatic activity dependent.

Many proteins are being regulated dynamically by methylation and demethylation, and the interplay of the methylation-demethylation machinery controls various processes such as gene expression, protein function, modification, and degradation ^48^. Therefore, it is possible that PGC-1α degradation could also be dynamically regulated by lysine methylation-demethylation cycle. We postulated that inhibiting lysine demethylation might facilitate PGC-1α degradation, an effect similar or comparable to SMYD5 overexpression-mediated methylation. To test this hypothesis, HEK293T cells were co-transfected with Flag-PGC-1α, along with HA alone vector or HA-SMYD5. In some groups, cells were only transfected with Flag-PGC-1α but treated with lysine-specific demethylase 1 (LSD1) inhibitor, pargyline ^49^. The result showed that exposure to pargyline led to a comparable reduction of PGC-1α half-life as did SMYD5 overexpression (**Figure 6D**), suggesting a dynamic control of PGC-1α stability by lysine methylation-demethylation balance. In all, these results clearly indicate that methylation of PGC-1α by SMYD5 leads to increased ubiquitination and consequently promotes proteasomal degradation of PGC-1α.

### Methyl-binding Protein PHF20L1 Is Involved in SMYD5-mediated PGC-1α Degradation

We have shown so far that SMYD5-mediated PGC-1α methylation triggers ubiquitin-dependent PGC-1α proteasomal degradation. However, it remains unclear how SMYD5-catalyzed PGC-1α methylation leads to its accelerated degradation. It has been reported that methylated lysine residues in substrate proteins interact with certain methyl-binding domain containing “readers”, which subsequently recruit, directly or indirectly, specific E3 ubiquitin ligases to regulate protein stability and turnover ^38^. We speculated that inhibiting the methyl lysine readers may block the binding/recognition of methylated lysine residues of PGC-1α by methyl lysine readers, thus preventing E3 ubiquitin ligase-mediated PGC-1α degradation. Previous study reported that the mono-methylated lysine 20 of histone 4 (H4K20) catalyzed by SMYD2 was recognized by the methyl-lysine reader L3MBTL1 (lethal 3 malignant brain tumor-like protein 1), a protein containing the malignant-brain-tumor (MBT) methylation-binding domain ^50^. UNC1215, a potent methyl-lysine binding protein inhibitor, has been shown to inhibit the cognate reader L3MBTL3 for binding to mono- or dimethyl lysine-containing peptides ^51^. However, knockdown of L3MBTL3 did not cause any significant change of PGC-1α protein level (**Supplementary Figure 7**).

It has been demonstrated that UNC1215, at high concentrations, inhibited other methyl-binding proteins, such as plant homeodomain finger protein 20-like 1 (PHF20L1), a protein that also contains the MBT domain ^52^. We treated HEK293T and HCT116 cells with UNC1215 and observed a significantly elevated expression of PGC-1α protein (**Figure 7A**) indicating that PGC-1α degradation is mediated, at least partially, by the methyl-lysine reader PHF20L1.

**Figure 7.**
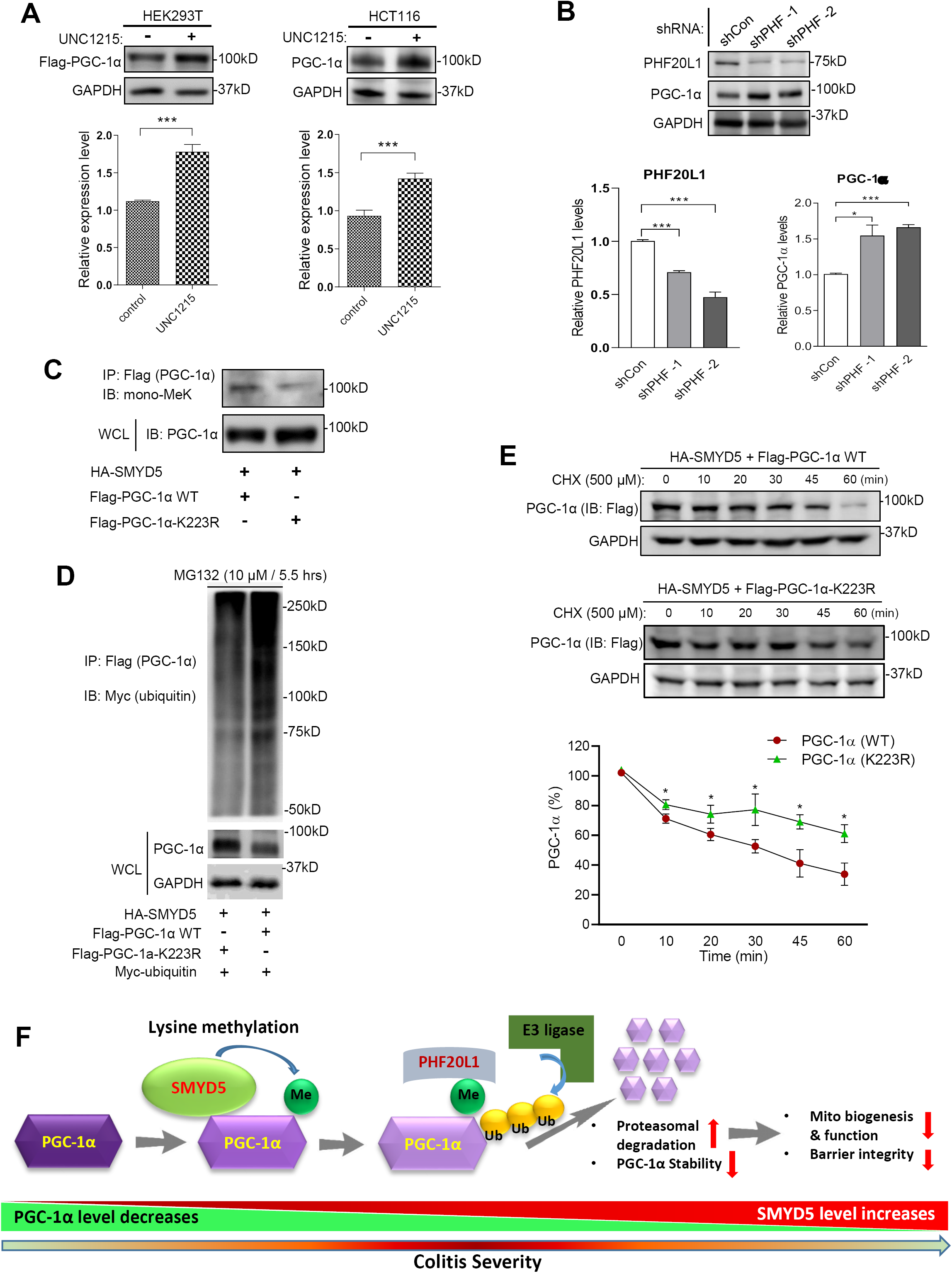
Methyl-binding protein PHF20L1 is involved in SMYD5-mediated PGC-1α degradation. (A) HEK293T cells overexpressing Flag-PGC-1α and HCT116 cells were treated with vehicle control or the methyl-binding protein inhibitor, UNC1215 (80 μM) for 24 hrs, and the whole cell lysates were immunoblotted for the expression of PGC-1α (anti-Flag for HEK293T) and GAPDH. ^***^*p* < 0.001, n = 3. (B) HCT116 cells were transduced with shRNA viruses targeting the methyl-binding protein PHF20L1 (shPHF −1 and shPHF −2) or control shRNA viruses (shCon), and whole cell lysates were immunoblotted for PHF20L1, PGC-1α, and GAPDH. ^*^*p* < 0.05; ^***^*p* < 0.001; n = 3. (C) HEK293T cells were transfected with HA-SMYD5 along with Flag-PGC-1α-WT or Flag-PGC-1α-K223R. Then whole cell lysates were subjected to immunoprecipitation with anti-Flag antibody (for Flag-PGC-1α WT or K223R) and the immunoprecipitated Flag-PGC-1α (WT or K223R) was analyzed by Western blot for lysine methylation using anti-mono methyl lysine (mono-MeK) antibody. Whole cell lysates (WCL) were also immunoblotted for PGC-1α (anti-Flag). (D) HEK293T cells were co-transfected with myc-ubiquitin, HA-SMYD5, and Flag-PGC-1α WT or Flag-PGC-1α-K223R followed by treatment with MG132 (10 μM) for 5.5 hrs. Then whole cell lysates were subjected to immunoprecipitation with anti-Flag antibody (for Flag-PGC-1α WT or K223R) and the immunoprecipitated Flag-PGC-1α (WT or K223R) was analyzed by Western blot for ubiquitination using anti-myc antibody (for myc-ubiquitin). Whole cell lysates (WCL) were also immunoblotted for PGC-1α (anti-Flag) and GAPDH. (E) HEK293T cells were transfected with HA-SMYD5 along with Flag-PGC-1α-WT or Flag-PGC-1α-K223R. Then cells were treated with cycloheximide (CHX, 500 μM) for indicated time periods before whole cell lysates were immunoblotted for PGC-1α (anti-Flag IgG) and GAPDH. The relative protein levels of Flag-PGC-1α WT or K223R at each time point (normalized to GAPDH) is plotted as a percentage (%) of the amount at 0 min. Flag-PGC-1α-WT vs. Flag-PGC-1α-K223R, ^*^*p* < 0.05; n = 3. (F) A schematic model depicting the SMYD5-catalyzed methylation, ubiquitination and degradation of PGC-1α protein in regulating intestinal health and colitis.

Previous study has shown that PHF20L1 binds to retinoblastoma tumor suppressor protein (pRB) at mono-methylated lysine 810 (K810) and recruits specific acetyltransferase complex to target genes ^53^. Interestingly, SMYD2 has been reported to mediate mono-methylation of pRB at K810 ^54^. It has been demonstrated that certain acetylated lysine residues target substrate proteins for ubiquitination and degradation ^55^. Intriguingly, it has been shown that PGC-1α is highly acetylated and ubiquitinated within the intestinal epithelium in mice after DSS exposure ^10^. Therefore, it is possible that mono-methylated K223 of PGC-1α may serve as a docking site for “readers” like PHF20L1 which recruits distinct acetyltransferase to “tag” PGC-1α for ubiquitination and degradation. To test this hypothesis, we generated clones of HCT116 cells that stably express shRNA targeting PHF20L1 (left panel, **Figure 7B**). Immunoblot analysis showed that PHF20L1 silencing substantially increased the expression of PGC-1α (right panel, **Figure 7B**), indicating that PHF20L1 may mediate PGC-1α lysine ubiquitination and its proteasomal degradation.

We next explored the role that lysine residue K223 plays in the process of PHF20L1-mediated PGC-1α ubiquitination and degradation. First, we constructed K223R point mutation in PGC-1α (Flag-PGC-1α-K223R) and co-expressed SMYD5 and wildtype PGC-1α (Flag-PGC-1α WT) or Flag-PGC-1α-K223R in HEK293T cells. Immunoblot analysis revealed that K223R mutation of PGC-1α significantly reduced SMYD5-mediated PGC-1α methylation and increased PGC-1α protein level (**Figure 7C**). This finding revealed a critical role of K223 mono-methylation in PGC-1α stability. In order to see whether K223R mutation would render PGC-1α to be less ubiquitinated due to failure of methylation of K223R mutant, we co-expressed myc-ubiquitin, HA-SMYD5, and Flag-PGC-1α WT or Flag-PGC-1α-K223R in HEK293T cells and performed co-immunoprecipitation assay. The result showed that K223R mutation indeed led to reduced PGC-1α ubiquitination (**Figure 7D**, top) and elevated PGC-1α protein level (**Figure 7D**, lower band), suggesting that K223 residue plays an important role in mediating PGC-1α ubiquitination. Furthermore, cycloheximide chase assay revealed a significantly increased half-life of PGC-1α K223R mutant as compared to PGC-1α WT (**Figure 7E**), suggesting that K223R mutation, at least partially, protects PGC-1α from ubiquitin-dependent proteasomal degradation.

Altogether, these results clearly demonstrated that SMYD5-catalyzed PGC-1α methylation at K223 residue, probably via methyl-binding protein PHF20L1, promotes PGC-1α ubiquitination and proteasomal degradation.

## Discussion

An emerging body of literature suggests that mitochondrial dysfunction is implicated in the pathogenesis and progression of IBD ^23^. It has been well established that regulation of mitochondrial functions requires PGC-1α, a transcriptional co-activator that induces mitochondrial biogenesis and antioxidant activity through interactions with multiple transcription factors, including NRF-1 or 2, which then stimulate the expression of TFAM and other mitochondrial matrix proteins essential for the replication and transcription of mtDNA ^56^. Recent study has demonstrated that decreased PGC-1α in IECs resulted in mitochondrial dysfunction during colitis^10^. However, the underlying molecular mechanism has not been fully elucidated.

In this study, we have provided the first evidence that SMYD5, which was initially identified as underlying molecular mechanism has not been fully elucidated. an epigenetic modifier that tri-methylates H4K20 to regulate downstream inflammatory genes of TLR4 in macrophages ^14^, functions as a critical negative regulator of epithelial integrity and intestinal homeostasis by suppressing mitochondrial biogenesis and function through methylation-mediated degradation of transcriptional coactivator PGC-1α (**Figure 7F**). We demonstrated that SYMD5 is expressed in IECs, whose expression was up-regulated in intestinal mucosa from IBD patients and DSS-induced colitic mice. SMYD5 knockdown in human IECs resulted in increased mitochondrial biogenesis and function and reduced ROS production. Further, IEC-specific SMYD5 deficiency protected mice against DSS-induced colitis and intestinal inflammation. Mechanistically, we observed that SMYD5 interacts directly and methylates PGC-1α, and methylation of PGC-1α in IECs leads to its ubiquitination and proteasome-mediated degradation. The expression of SMYD5 in intestinal epithelia, along with its ability to interact with and methylate PGC-1α, suggests that SMYD5 and PGC-1α may work together to regulate mitochondrial function and intestinal homeostasis.

Although members of the SMYD family have been studied for decades, the study of SMYD5 is still very limited. Our finding that SMYD5 is upregulated in the colon samples from IBD patients is consistent with a previous study that, by using GWAS, has demonstrated the upregulation of SMYD5 in human IBD colonic biopsies ^57^. Recent proteomic studies have showed that compared with other members of SMYD family, SMYD5 is broadly expressed, but with a medium level in colon and rectum in healthy tissues ^58^. Meanwhile, other studies have shown that in cancer patients, especially in patients with gastrointestinal cancers, SMYD5 transcription is significantly up-regulated compared with that from healthy individuals ^59^. It is well documented that IBD is the primary risk factor for the development of gastrointestinal cancers ^60^. Together, this suggests that SMYD5 may play a critical role in intestinal epithelia during the pathogenesis of various gastrointestinal diseases.

It is possible that SMYD5 may exert its role in the intestinal epithelium through other mechanisms as well. It has been reported that SMYD5 depletion leads to increased cell growth due to decreased genome-wide H4K20 tri-methylation ^15^. In our study, upon DSS administration, IECs from Smyd5^ΔIEC^ mice displayed increased cell proliferation compared with those from Smyd5^fl/fl^ mice (data not shown). This suggests that SMYD5 may also play a role in the nucleus of IECs during the pathogenesis of IBD, probably independent of PGC-1α.

We have also identified a lysine residue, K223 of murine PGC-1α, that was mono-methylated by SMYD5, and we further mutated this lysine residue to arginine (K223R). This point mutation has significantly reduced SMYD5-mediated PGC-1α methylation and ubiquitination. More importantly, K223R mutation has increased the half-life of PGC-1α. These findings have supported the notion that SMYD5-mediated PGC-1α methylation is involved in regulating PGC-1α turnover in IECs. Interestingly, a recent study also revealed the mono-methylation of human PGC-1α at K224 (equivalent to the K223 of mouse PGC-1α) induced by hypoxia in human glioblastoma cells and PGC-1α K224 demonstrated limited mono-methylation under normoxic condition ^61^. However, they did not detect any changes of PGC-1α mRNA or protein expression. Instead, they found that hypoxia decreases PGC-1α activity via inhibition of the lysine demethylase 3A (KDM3A)-mediated de-methylation of human PGC-1α at K224, which consequently results in reduced mitochondrial biogenesis ^61^. They also observed that treatment of human glioblastoma cells with 5-carboxy-8-hydroxyquinoline (IOX1), a potent broad-spectrum inhibitor of the JMJD family of 2-oxoglutaratedependent demethylases, or deferoxamine (DFO), an iron chelator that blocks iron-dependent demethylation, drastically increased PGC-1α K224 monomethylation, suggesting that PGC-1α K224 monomethylation induced by hypoxia is due to inhibition of a demethylase. However, in our present study, treatment of IECs with either IOX1 or DFO failed to induce any change in PGC-1α methylation or degradation (data not shown), suggesting that the mechanism and effect of PGC-1α methylation on mitochondrial biogenesis and function may be cell type or context dependent.

In our study, immunoblot analysis has revealed methylated lysine signals at N-terminal, middle, and C-terminal (weak signal) regions of PGC-1α. However, our MS analysis only detected a lysine methylation at K223 within the middle fragment of PGC-1α (aa 190-345). We speculate this may be due to technique challenge. Interestingly, the N-terminal domain of PGC-1α has been reported to have no effect on protein stability and subcellular distribution ^43^. Recently, a lysine residue within the C-terminal (K779) of PGC-1α has been reported to be monomethylated by SET7/9 in Hepa 1-6 cells; however, this methylation has no effect on PGC-1α protein stability ^62^. Nonetheless, methylation on other lysine residues of PGC-1α may also contribute to its ubiquitination and degradation. Of note, K223R mutation failed to completely reverse SMYD5 mediated PGC-1α methylation and degradation. Therefore, it is possible that SMYD5 catalyze methylation of multiple lysine residues within PGC-1α, which collectively contribute to PGC-1α degradation, though K223 methylation may play the major role. It will be interesting to identify all the lysine residues in PGC-1α methylated by SMYD5 and define their distinct functional consequences.

It has been reported that the MYND domain within the SMYD proteins are involved in interactions with a preference for binding to a proline-rich motif (PXLXP) ^63^. For instance, SMYD1 has been reported to interact with PPLIP motif of skeletal and heart muscle-specific variant of the alpha subunit of nascent polypeptide associated complex ^63^. Moreover, the PXL motif in HSP90 and P23 has been reported to mediate their binding with SMYD2 ^64^. Interestingly, PGC-1α also contains a proline-rich region ^65^. Therefore, it is possible that the proline-rich motif of PGC-1α may mediate its interaction with SMYD5, which warrants further investigation.

In summary, our present study, for the first time, revealed a protective role of SMYD5 ablation in IECs during IBD pathogenesis and the underlying mechanism. We identified the first non-histone substrate of SMYD5, PGC-1α, the master regulator of mitochondrial functions. By controlling the methylation status of PGC-1α, SMYD5 modulates its stability and turnover, which further regulates the mitochondria function in IECs. Limited research has been done on SMYD5, with focus solely on histone biology and epigenetics. Our findings have broadened the understanding of the members of SMYD family, and also provided the evidence that targeting SMYD5/PGC-1α axis in IECs could be a potential therapeutic target for IBD treatment, as well as other mitochondria-involved disease conditions such as neurodegenerative diseases, cancers, obesity and diabetes.

## Acknowledgments and grant support

We thank Drs. Didier Merlin and Ming-Hui Zou for their intellectual input of the study, and Qian (Zoe) Liu for technical support for the mice. This work was supported by the National Institutes of Health grant number R01HL128647 (to C.L.), Georgia State University faculty start-up fund (to C.L.), and R01HL128014 (to Z.X.)

## Abbreviations used in this paper

CD: Crohn’s disease
CHX: cycloheximide
COX I/II: cytochrome c oxidase I/II
DAI: disease activity index
DSS: dextran sulfate sodium
FCCP: trifluoromethoxy carbonyl cyanide phenylhydrazone
FITC: fluorescein isothiocyanate
H4K20: histone 4 lysine 20
HA: hemagglutinin
HCT116: human colorectal carcinoma cell line
IBD: inflammatory bowel disease
IEC: intestinal epithelial cell
K223: lysine 223
K223R: lysine 223 to arginine
KO: knockout
L3MBTL1/3: lethal (3) malignant brain tumor-like protein 1/3
LPS: lipopolysaccharide
LSD1: lysine-specific demethylase 1
mtDNA: mitochondrial DNA
OCR: oxygen consumption rate
OE: overexpression
PHF20L1: plant homeodomain finger protein 20-like 1
PGC-1α: peroxisome proliferator-activated receptor gamma coactivator 1 alpha
SMYD5: SET and MYND domain- containing protein 5
TEM: transmission electron microscopy
Tfam/TFAM: mitochondrial transcription factor A
UC: ulcerative colitis

## Notes

**Disclosures:** The authors declare no conflict of interests.

### Competing Interest Statement

The authors have declared no competing interest.

## References

1. Abraham C, Cho JH. Inflammatory bowel disease. N Engl J Med 2009;361:2066–78.

2. Kappelman MD, Rifas-Shiman SL, Kleinman K, et al. The prevalence and geographic distribution of Crohn’s disease and ulcerative colitis in the United States. Clin Gastroenterol Hepatol 2007;5:1424–9.

3. Kappelman MD, Rifas-Shiman SL, Porter CQ, et al. Direct health care costs of Crohn’s disease and ulcerative colitis in US children and adults. Gastroenterology 2008;135:1907–13.

4. Peterson LW, Artis D. Intestinal epithelial cells: regulators of barrier function and immune homeostasis. Nat Rev Immunol 2014;14:141–53.

5. Ho GT, Aird RE, Liu B, et al. MDR1 deficiency impairs mitochondrial homeostasis and promotes intestinal inflammation. Mucosal Immunol 2018;11:120–130.

6. Jackson DN, Panopoulos M, Neumann WL, et al. Mitochondrial dysfunction during loss of prohibitin 1 triggers Paneth cell defects and ileitis. Gut 2020;69:1928–1938.

7. Cherry AD, Piantadosi CA. Regulation of mitochondrial biogenesis and its intersection with inflammatory responses. Antioxid Redox Signal 2015;22:965–76.

8. Wu Z, Puigserver P, Andersson U, et al. Mechanisms controlling mitochondrial biogenesis and respiration through the thermogenic coactivator PGC-1. Cell 1999;98:115–24.

9. D’Errico I, Salvatore L, Murzilli S, et al. Peroxisome proliferator-activated receptor-gamma coactivator 1-alpha (PGC1alpha) is a metabolic regulator of intestinal epithelial cell fate. Proc Natl Acad Sci U S A 2011;108:6603–8.

10. Cunningham KE, Vincent G, Sodhi CP, et al. Peroxisome Proliferator-activated Receptor-gamma Coactivator 1-alpha (PGC1alpha) Protects against Experimental Murine Colitis. J Biol Chem 2016;291:10184–200.

11. Ussakli CH, Ebaee A, Binkley J, et al. Mitochondria and tumor progression in ulcerative colitis. J Natl Cancer Inst 2013;105:1239–48.

12. Du SJ, Tan X, Zhang J. SMYD Proteins: Key Regulators in Skeletal and Cardiac Muscle Development and Function. Anat Rec (Hoboken) 2014;297:1650–62.

13. Spellmon N, Holcomb J, Trescott L, et al. Structure and Function of SET and MYND Domain-Containing Proteins. International Journal of Molecular Sciences 2015;16.

14. Stender JD, Pascual G, Liu W, et al. Control of proinflammatory gene programs by regulated trimethylation and demethylation of histone H4K20. Mol Cell 2012;48:28–38.

15. Kidder BL, Hu G, Cui K, et al. SMYD5 regulates H4K20me3-marked heterochromatin to safeguard ES cell self-renewal and prevent spurious differentiation. Epigenetics Chromatin 2017;10:8.

16. Fujii T, Tsunesumi S, Sagara H, et al. Smyd5 plays pivotal roles in both primitive and definitive hematopoiesis during zebrafish embryogenesis. Sci Rep 2016;6:29157.

17. Farooq SM, Hou Y, Li H, et al. Disruption of GPR35 Exacerbates Dextran Sulfate Sodium-Induced Colitis in Mice. Dig Dis Sci 2018;63:2910–2922.

18. Arora K, Sinha C, Zhang W, et al. Altered cGMP dynamics at the plasma membrane contribute to diarrhea in ulcerative colitis. Am J Pathol 2015;185:2790–804.

19. Nowarski R, Jackson R, Gagliani N, et al. Epithelial IL-18 Equilibrium Controls Barrier Function in Colitis. Cell 2015;163:1444–56.

20. Wu S, Lu Q, Ding Y, et al. Hyperglycemia-Driven Inhibition of AMP-Activated Protein Kinase alpha2 Induces Diabetic Cardiomyopathy by Promoting Mitochondria-Associated Endoplasmic Reticulum Membranes In Vivo. Circulation 2019;139:1913–1936.

21. Wu S, Lu Q, Wang Q, et al. Binding of FUN14 Domain Containing 1 With Inositol 1,4,5-Trisphosphate Receptor in Mitochondria-Associated Endoplasmic Reticulum Membranes Maintains Mitochondrial Dynamics and Function in Hearts in Vivo. Circulation 2017;136:2248–2266.

22. Nguyen HT, Dalmasso G, Torkvist L, et al. CD98 expression modulates intestinal homeostasis, inflammation, and colitis-associated cancer in mice. J Clin Invest 2011;121:1733–47.

23. Novak EA, Mollen KP. Mitochondrial dysfunction in inflammatory bowel disease. Front Cell Dev Biol 2015;3:62.

24. Okayasu I, Hatakeyama S, Yamada M, et al. A novel method in the induction of reliable experimental acute and chronic ulcerative colitis in mice. Gastroenterology 1990;98:694–702.

25. Doughan M, Spellmon N, Li C, et al. SMYD proteins in immunity: dawning of a new era. AIMS Biophys 2016;3:450–455.

26. Ito R, Shin-Ya M, Kishida T, et al. Interferon-gamma is causatively involved in experimental inflammatory bowel disease in mice. Clin Exp Immunol 2006;146:330–8.

27. Sanchez-Munoz F, Dominguez-Lopez A, Yamamoto-Furusho JK. Role of cytokines in inflammatory bowel disease. World J Gastroenterol 2008;14:4280–8.

28. Safdar A, Little JP, Stokl AJ, et al. Exercise increases mitochondrial PGC-1alpha content and promotes nuclear-mitochondrial cross-talk to coordinate mitochondrial biogenesis. J Biol Chem 2011;286:10605–17.

29. LeBleu VS, O’Connell JT, Gonzalez Herrera KN, et al. PGC-1alpha mediates mitochondrial biogenesis and oxidative phosphorylation in cancer cells to promote metastasis. Nat Cell Biol 2014;16:992-1003, 1-15.

30. Larsson NG, Wang J, Wilhelmsson H, et al. Mitochondrial transcription factor A is necessary for mtDNA maintenance and embryogenesis in mice. Nat Genet 1998;18:231–6.

31. Chicherin IV, Dashinimaev E, Baleva M, et al. Cytochrome c Oxidase on the Crossroads of Transcriptional Regulation and Bioenergetics. Front Physiol 2019;10:644.

32. Bar F, Bochmann W, Widok A, et al. Mitochondrial gene polymorphisms that protect mice from colitis. Gastroenterology 2013;145:1055-1063.e3.

33. Liu Y, Shim E, Crespo-Mejias Y, et al. Cardiomyocytes are Protected from Antiretroviral Nucleoside Analog-Induced Mitochondrial Toxicity by Overexpression of PGC-1alpha. Cardiovasc Toxicol 2015;15:224–31.

34. Antoni L, Nuding S, Wehkamp J, et al. Intestinal barrier in inflammatory bowel disease. World J Gastroenterol 2014;20:1165–79.

35. Kang T, Lu W, Xu W, et al. MicroRNA-27 (miR-27) targets prohibitin and impairs adipocyte differentiation and mitochondrial function in human adipose-derived stem cells. J Biol Chem 2013;288:34394–402.

36. Kontaki H, Talianidis I. Lysine methylation regulates E2F1-induced cell death. Mol Cell 2010;39:152–60.

37. Elkouris M, Kontaki H, Stavropoulos A, et al. SET9-Mediated Regulation of TGF-beta Signaling Links Protein Methylation to Pulmonary Fibrosis. Cell Rep 2016;15:2733–44.

38. Leng F, Yu J, Zhang C, et al. Methylated DNMT1 and E2F1 are targeted for proteolysis by L3MBTL3 and CRL4(DCAF5) ubiquitin ligase. Nat Commun 2018;9:1641.

39. Hsiao K, Zegzouti H, Goueli SA. Methyltransferase-Glo: a universal, bioluminescent and homogenous assay for monitoring all classes of methyltransferases. Epigenomics 2016;8:321–39.

40. Wilkinson AW, Diep J, Dai S, et al. SETD3 is an actin histidine methyltransferase that prevents primary dystocia. Nature 2019;565:372–376.

41. Zhang X, Tanaka K, Yan J, et al. Regulation of estrogen receptor alpha by histone methyltransferase SMYD2-mediated protein methylation. Proc Natl Acad Sci U S A 2013;110:17284–9.

42. Olson BL, Hock MB, Ekholm-Reed S, et al. SCFCdc4 acts antagonistically to the PGC-1alpha transcriptional coactivator by targeting it for ubiquitin-mediated proteolysis. Genes Dev 2008;22:252–64.

43. Trausch-Azar J, Leone TC, Kelly DP, et al. Ubiquitin proteasome-dependent degradation of the transcriptional coactivator PGC-1{alpha} via the N-terminal pathway. J Biol Chem 2010;285:40192–200.

44. Wei P, Pan D, Mao C, et al. RNF34 is a cold-regulated E3 ubiquitin ligase for PGC-1alpha and modulates brown fat cell metabolism. Mol Cell Biol 2012;32:266–75.

45. Park JH, Kang HJ, Lee YK, et al. Inactivation of EWS reduces PGC-1alpha protein stability and mitochondrial homeostasis. Proc Natl Acad Sci U S A 2015;112:6074–9.

46. Hsin IL, Sheu GT, Jan MS, et al. Inhibition of lysosome degradation on autophagosome formation and responses to GMI, an immunomodulatory protein from Ganoderma microsporum. Br J Pharmacol 2012;167:1287–300.

47. Lecker SH, Goldberg AL, Mitch WE. Protein degradation by the ubiquitin-proteasome pathway in normal and disease states. J Am Soc Nephrol 2006;17:1807–19.

48. Chakraborty A, Viswanathan P. Methylation-Demethylation Dynamics: Implications of Changes in Acute Kidney Injury. Anal Cell Pathol (Amst) 2018;2018:8764384.

49. Shi Y, Lan F, Matson C, et al. Histone demethylation mediated by the nuclear amine oxidase homolog LSD1. Cell 2004;119:941–53.

50. Boehm D, Jeng M, Camus G, et al. SMYD2-Mediated Histone Methylation Contributes to HIV-1 Latency. Cell Host Microbe 2017;21:569–579 e6.

51. James LI, Barsyte-Lovejoy D, Zhong N, et al. Discovery of a chemical probe for the L3MBTL3 methyllysine reader domain. Nat Chem Biol 2013;9:184–91.

52. Esteve PO, Terragni J, Deepti K, et al. Methyllysine reader plant homeodomain (PHD) finger protein 20-like 1 (PHF20L1) antagonizes DNA (cytosine-5) methyltransferase 1 (DNMT1) proteasomal degradation. J Biol Chem 2014;289:8277–87.

53. Carr SM, Munro S, Sagum CA, et al. Tudor-domain protein PHF20L1 reads lysine methylated retinoblastoma tumour suppressor protein. Cell Death Differ 2017;24:2139–2149.

54. Cho HS, Hayami S, Toyokawa G, et al. RB1 methylation by SMYD2 enhances cell cycle progression through an increase of RB1 phosphorylation. Neoplasia 2012;14:476–86.

55. Narita T, Weinert BT, Choudhary C. Functions and mechanisms of non-histone protein acetylation. Nat Rev Mol Cell Biol 2019;20:156–174.

56. Luo C, Widlund HR, Puigserver P. PGC-1 Coactivators: Shepherding the Mitochondrial Biogenesis of Tumors. Trends Cancer 2016;2:619–631.

57. Wu F, Dassopoulos T, Cope L, et al. Genome-wide gene expression differences in Crohn’s disease and ulcerative colitis from endoscopic pinch biopsies: insights into distinctive pathogenesis. Inflamm Bowel Dis 2007;13:807–21.

58. Kim MS, Pinto SM, Getnet D, et al. A draft map of the human proteome. Nature 2014;509:575–81.

59. Song J, Liu Y, Chen Q, et al. Expression patterns and the prognostic value of the SMYD family members in human breast carcinoma using integrative bioinformatics analysis. Oncol Lett 2019;17:3851–3861.

60. Kim ER, Chang DK. Colorectal cancer in inflammatory bowel disease: the risk, pathogenesis, prevention and diagnosis. World J Gastroenterol 2014;20:9872–81.

61. Qian X, Li X, Shi Z, et al. KDM3A Senses Oxygen Availability to Regulate PGC-1alpha-Mediated Mitochondrial Biogenesis. Mol Cell 2019;76:885–895 e7.

62. Aguilo F, Li S, Balasubramaniyan N, et al. Deposition of 5-Methylcytosine on Enhancer RNAs Enables the Coactivator Function of PGC-1alpha. Cell Rep 2016;14:479–92.

63. Sims RJ, 3rd, Weihe EK, Zhu L, et al. m-Bop, a repressor protein essential for cardiogenesis, interacts with skNAC, a heart- and muscle-specific transcription factor. J Biol Chem 2002;277:26524–9.

64. Obermann WMJ. A motif in HSP90 and P23 that links molecular chaperones to efficient estrogen receptor alpha methylation by the lysine methyltransferase SMYD2. J Biol Chem 2018;293:16479–16487.

65. Vercauteren K, Gleyzer N, Scarpulla RC. PGC-1-related coactivator complexes with HCF-1 and NRF-2beta in mediating NRF-2(GABP)-dependent respiratory gene expression. J Biol Chem 2008;283:12102–11.

