## Supplementary Methods and Figures for "Methyltransferase SMYD5 Exaggerates IBD by Downregulating Mitochondrial Functions via Post-translational Control of PGC-1α Stability"

**Short title:** Up-regulated SMYD5 inhibits PGC-1α stability in IBD


**Authors:**

Yuning Hou [1], Xiaonan Sun [1], Pooneh Tavakoley Gheinani [1], Xiaoqing Guan [1], Shaligram Sharma [1], Yu Zhou [1,2], Chengliu Jin [3], Zhe Yang [4], Anjaparavanda P. Naren [5], Jun Yin [6], Timothy L. Denning [7], Andrew T. Gewirtz [7], Zhonglin Xie [1], Chunying Li [1,*]

[1] Center for Molecular and Translational Medicine, Georgia State University, Atlanta, GA 30303.

[2] Division of Vascular Surgery, The First Affiliated Hospital, Sun Yat-Sen University, Guangzhou, China

[3] Transgenic and Gene Targeting Core, Georgia State University, Atlanta, GA 30303.

[4] Department of Biochemistry, Microbiology, and Immunology, Wayne State University School of Medicine, Detroit, MI 48201.

[5] Department of Pediatrics, Division of Pulmonary Medicine, Cincinnati Children's Hospital Medical Center, Cincinnati, OH 45229.

[6] Department of Chemistry, Center for Diagnostics & Therapeutics, Georgia State University, Atlanta, GA 30303.

[7] Center for Inflammation, Immunity and Infection, Institute for Biomedical Sciences, Georgia State University, Atlanta, GA 30303.

**Correspondence:**

Chunying Li, Ph.D.

Room 511 Research Science Center, 157 Decatur St SE, Atlanta, GA 30303.

### Supplementary Methods

#### *Quantitative Reverse-transcription Polymerase Chain Reaction (RT-qPCR) Analysis*

Total RNA was extracted from cells with TRIzol reagent (AM9738, Invitrogen) and cDNA was synthesized using a Denville rAMP™ cDNA Synthesis kit (CC1151, Denville) according to the manufacturer's instructions. Quantitative reverse-transcriptase polymerase chain reaction (RT-qPCR) analysis was performed to determine the expression levels of mitochondrial biogenesis markers (Tfam, Cox I, and Cox II) in human IEC cell lines and mouse IECs. RT-PCR was performed in triplicate using power up SYBR Green master mix (A25741, Applied Biosystems). The expression levels of Tfam, Cox I, and Cox II were normalized to the level of GAPDH mRNA. The respective primers used are listed in **Supplementary Table 2**.

#### *Detection of Mitochondrial DNA Copy Number*

Total DNA was extracted from cells using Blood & Cell Culture DNA Mini Kit (13323, Invitrogen) according to the manufacturer's instructions. Mitochondrial DNA copy number was determined by RT-qPCR as previously reported [1]. In general, the mitochondrial gene (NADH dehydrogenase) and nuclear gene (β-actin) were determined and the respective primers used are listed in **Supplementary Table 2**. The results are presented as ratio of mitochondrial DNA relative to nuclear DNA.

#### *Cell Culture, Transfection, and Treatment*

Human colorectal epithelial cells (HCT116, HT-29, and Caco-2) and HEK293T cells were cultured under standard culture conditions as described before [2, 3]. In brief, colorectal epithelial cells and HEK293 cells were cultured in MaCoy's 5A (SH30200.1, Hyclone) or Dulbecco's modified Eagle's medium (DMEM, 11965092, Gibco) medium supplemented with 10% fetal bovine serum (FBS) and antibiotics (100 U/ml penicillin and streptomycin). The cells were maintained in 37 °C in an incubator with 5% $CO_2$. When the confluence reaches 80%, cells were used for transfection. For transient transfection, plasmids encoding wildtype SMYD5 (pcDNA3.1-HA-SMYD5) (Genscript Biotech), enzymatically inactive SMYD5 (pcDNA3.1-HA-SMYD5-H316L), wildtype PGC-1α (pcDNA-Flag-PGC-1α) (#1026, Addgene) or methylation resistant PGC-1α (pcDNA-Flag-PGC-1α K223R), ubiquitin (pcDNA3.1-Myc-ubiquitin), or empty vector plasmid were transfected into cells using Lipofectamine LTX according to the manufacturer's instruction.

To generate methylation resistant PGC-1α, pcDNA-Flag-PGC-1α, which expressing Flag-tagged wildtype PGC-1α, was used as template. Q5® Site-Directed Mutagenesis Kit (E0554S,

New England Biolabs) was used to generate K223R mutation using primers synthesized by Sigma-Aldrich according to the manufacturer's instruction.

For evaluating cellular oxidative stress induced by inflammatory response and subcellular distribution of PGC-1α and SMYD5 upon exposure to proinflammatory cytokines, cells were treated with or without TNF-α (Prospec) (20 ng/ml) and/or IFN-γ (Prospec) (20 ng/ml) for 24 hrs. Hydrogen peroxide ($H_2O_2$) was used to treat the cells to define the role of SMYD5 in oxidative stress induced cell apoptosis. Briefly, cells were treated with 100 µM $H_2O_2$ for 0, 4, or 8 hrs, respectively. Then, cells were subjected to Western blot analysis to evaluate the cleavage of caspases. In order to investigate the pathways for PGC-1α degradation, cells were treated with proteasomal inhibitor (MG132, 10 µM, for 0, 16, and 24 hrs) or autophagy inhibitors (Bafilomycin A1 (10 nM) and chloroquine (50 µM) for 0, 8, 16, and 24 hrs) respectively followed by Western blotting. To test the role of SMYD5 on PGC-1α ubiquitination, cells were treated with MG132 (10 µM) for 5.5 hrs and were then subjected to immunoblotting analysis. For the purpose of investigating the importance of methyltransferase activity of SMYD5 on the proteasomal degradation of PGC-1α, cells were treated with or without lysine demethylase inhibitor (pargyline, 2.5 mM) for 24 hrs and then were treated with protein synthesis inhibitor (cycloheximide, CHX, 500 µM) for 0, 10, 20, 30, 45, and 60 min, respectively. Then, the cells were used for Western blot analysis. In order to determine the role of PHF20L1 in SMYD5-mediated PGC-1α degradation, cells were incubated with the antagonist of methyl-lysine reader (UNC1215, 80 µM, 24 hrs) followed by Western blotting to detect the expression of PGC-1α. Iron-dependent histone lysine demethylase inhibitor Desferrioxamine (DFO, 500 µM) and JmjC histone demethylase inhibitor (IOX1, 500 µM) were used to treat the cells (for 12 hrs) to investigate the effects of demethylase on PGC-1α expression.

#### *Histology, Immunofluorescence, and Immunochemistry*

For histological analysis, the "Swiss rolls" method was used to prepare the tissues as we reported before [2]. Then the tissues were fixed overnight in 4% paraformaldehyde, embedded in paraffin, and cut into 5-µm sections.

For immunofluorescence and immunohistochemistry analysis, antigen retrieval was performed by heat-induced epitope retrieval method using sodium citrate buffer (pH 6.0) in a water bath. Endogenous peroxidase was inhibited by blocking solution (SP-6000, Vector Laboratories) according to the manufacturer's instruction. The sections were blocked for 1 hour with TBST containing 5% normal goat serum (G9023, Sigma). The sections were incubated with the following primary antibodies overnight at 4˚C: anti-SMYD5 (NBP1-31222, Novus), 1:100 for immunohistochemistry; anti-SMYD5 (ab81419, Abcam), 1:100 for immunofluorescence; anti-PGC-1α (NBP1-04676, Novus), 1:150; and anti-mitochondria antibody (MAB1273, Millipore), 1:50. Followed by incubation with primary antibodies, the sections were washed with TBST three times and incubated for 1 hour at room temperature with Alexa Fluor 568 goat anti-rabbit IgG

(H+L) (A11036, Invitrogen) or Alexa Fluor 488 goat anti rabbit IgG (H+L) (A11034, Invitrogen). The sections were then washed, stained with DAPI for 5 min, and covered with antifade mountant (P36970, Invitrogen).

For immunohistochemistry staining, goat anti-rabbit IgG (H+L)-peroxidase conjugated secondary antibody (A0545, Invitrogen) was used. Then the sections were washed and incubated with peroxidase substrate (SK-4105, Vector Laboratories) for 10 min. Sections were then washed with water to remove extra substrate, processed for dehydration, and covered with Permount™ Mounting Medium (SP-15, Fisher scientific). The degree of staining was estimated with Image J by experienced pathologists and technicians who were blinded to the study.

#### *Western Blot Analysis*

Western blot analysis was performed according to standard procedures using lysates from tissues or cells as described before [2, 4] . In brief, tissues or cells were solubilized in M-PER mammalian protein extraction reagent (78501, Thermo Scientific) supplemented with protease inhibitor cocktail (P8340, Sigma). After the protein concentration was measured with Bradford method, same amounts of total proteins were separated by sodium dodecyl sulfate polyacrylamide gel, and transferred to polyvinylidene difluoride membranes (#1620177, Bio-Rad). The membranes were blocked with 5% goat serum at room temperature for 1 hour before the addition of primary antibodies for overnight incubation at 4˚C. The following day, the membranes were incubated with secondary antibodies for 1 h at room temperature. The ECL chemiluminescent kit was used for development, and the images were captured using Amersham Imager 600 RGB system (GE health care). The primary antibodies used were as follows: anti-SMYD5 (ab81419, Abcam), anti-GAPDH (AM4300, Invitrogen), anti-ubiquitin (#13-1600, Invitrogen), anti-PHF20L1 (NBP1-79401), anti-PGC-1α (NBP1-04676, Novus), anti-COX I (sc-19998, Santa Cruz), anti-COX II (#12282, Cell Signaling Technology), anti-COX IV (#4850, Cell Signaling Technology), anti-HSP60 (#12165, Cell Signaling Technology), anti-Pyruvate Dehydrogenase (#3205, Cell Signaling Technology), anti-SDHA (#11998, Cell Signaling Technology), anti-VDAC (#4661, Cell Signaling Technology), anti-UCP2 (#89326, Cell Signaling Technology), anti-UCP3 (#14670, Cell Signaling Technology), anti-cleaved caspase 3 (#9664, Cell Signaling Technology), anti-cleaved caspase 9 (#7237, Cell Signaling Technology), anti-TFAM (ABE483, Millipore), anti-myc (#05-724, Millipore), anti-Flag (F9291, Sigma), anti-HA (H9658, Sigma).

#### *Immunoprecipitation*

Immunoprecipitation assay was used for detecting the methylation status of lysine residues in PGC-1α. Briefly, HEK293T cells were co-transfected with HA-tagged SMYD5 and wild-type Flag-tagged PGC-1α or co-transfected with HA-tagged SMYD5 and mutated Flag-tagged PGC-1α (K223R). Forty-eight hours post-transfection, cells were lysed with M-PER mammalian protein extraction reagent supplemented with protease inhibitor cocktail, and the cleared lysates were incubated with anti-Flag Magnetic beads (M8823, Sigma) at 4˚C overnight. After washing, the PGC-1α was eluted by boiling in Laemmli sample buffer. Immunoblot analysis was conducted to detect the mono-methylated lysine (#16479, Cell Signaling Technology), di-methylated lysine (#14117, Cell Signaling Technology), and tri-methylated lysine (#14680, Cell Signaling Technology), as described above.

***Co-immunoprecipitation***

Co-immunoprecipitation assay was used for detecting the interaction between SMYD5 and PGC-1α, and detecting the ubiquitination of PGC-1α. Generally, HCT116 and HEK293T cells were co-transfected with Flag-tagged PGC-1α and myc-tagged ubiquitination or co-transfected with Flag-tagged PGC-1α and HA-tagged SMYD5. After 48 hrs of culture, the cells were solubilized in M-PER mammalian protein extraction reagent supplemented with protease inhibitor cocktail, and the cleared lysates were incubated with anti-Flag magnetic beads (M8823, Sigma) or anti-HA-Agarose (A2095, Sigma) at 4˚C overnight. After extensive washing to remove unbound proteins, the bead-bound proteins were eluted by boiling in Laemmli sample buffer, followed by immunoblot analysis as described above.

***Transmission Electron Microscopy***

Sample embedding, sectioning, and transmission electron microscopy (TEM) were performed by the Emory University Robert P. Apkarian Integrated Electron Microscopy Core as described before [5]. Colon tissues obtained from Smyd5$^{fl/fl}$ mice and Smyd5$^{\Delta IEC}$ mice were fixed with 2.5% glutaraldehyde in 0.1 mol/L cacodylate buffer (pH 7.4) followed by post-fixation with 1% osmium and 1.5% potassium ferrocyanide in the same buffer. Tissues were then dehydrated in ethanol and embedded in Eponate 12 resin. Ultrathin (70 nm) sections were cut with an ultramicrotome and stained with 5% uranyl acetate and 2% lead citrate. Images were acquired using a Hitachi H-7500 transmission electron microscope equipped with a SIA L12C 16 megapixel CCD camera. All measurement analyses were performed by laboratory personnel blinded to sample identities using ImageJ software (NIH). The Robert P. Apkarian Integrated Electron Microscopy Core (IEMC) at Emory University is subsidized by the School of Medicine and Emory College of Arts and Sciences. Additional support was provided by the Georgia Clinical & Translational Science Alliance of the National Institutes of Health under award number UL1TR000454. The content is solely the responsibility of the authors and does not necessarily reflect the official views of the National Institutes of Health.

***Cellular oxidative stress measured using a fluorescence-based assay***

The cellular oxidative stress was measured using CellROX$^{TM}$ Orange Reagent (C10443, Fisher) as described in previous study [6] without or with exposure to proinflammatory cytokines (TNF-α and IFN-γ) in SMYD5 KO HCT116 cells and parental HCT116 cells (SMYD5 WT). The cells were treated as described above.

#### *Mass Spectrometry*

To identify the SMYD5 methylated lysine residues in PGC-1α, liquid chromatography with tandem mass spectrometry (LC-MS/MS) was used. Briefly, methylated PGC-1α were subjected to electrophoresis in SDS-PAGE gel and stained with 0.5% Coomassie Blue R-250. The gel was stored in 5% acetic acid and were subsequently processed and analyzed by System Mass Spectrometry Core Facility in Georgia Institute of Technology as described previously [7].

#### *Methyltranferase-Glo Assay*

In order to study the methylation status of lysine in PGC-1α, MTase-Glo assays were performed by multistep format in 96-well plate using MTase-Glo methayltransferase assay kit (V7601, Promega) according to the manufacturer's instruction [8]. Briefly, GST tagged PGC-1α and GST tagged SMYD5 were added into 20 µl reaction buffer (20 mM Tris, pH 8.0, 50 mM NaCl, 1 mM EDTA, 3 mM $MgCl_2$, and 0.1 mg/ml BSA) containing SAM (10 µM final concentration) and incubated for 30 min at room temperature. After the incubation, 5 ul MTase-Glo Reagent provided in the kit were added and incubated for 30 min at room temperature. Once the incubation was completed, 25 µl MTase-Glo Detection solution was added and incubated for 30 min at room temperature. Afterwards, the luminescence was measured with a plate-reading luminometer. In control group, identical reaction conditions were applied except purified GST protein was used instead of GST tagged SMYD5. For all experiments, all reactions were done in triplicate.

#### *In vitro methyltransferase assay of lysine methylation of PGC-1α*

Bacterially purified GST-SMYD5 or GST alone was mixed with GST-PGC-1α fragments (GST-PGC-1α aa 1-190, GST-PGC-1α aa 190-345, and GST-PGC-1α aa 345-797) in a methylation reaction as in ***Methyltranferase-Glo Assay***. Then, the reaction mixtures were subjected to Western blotting using a methyl lysine-specific antibody (ab23366, Abcam; Cat# 14117, Cat# 14679, Cell Signaling) to detect methylated lysine(s) of PGC-1α fragments.

### Supplementary References:


1. Zheng P, Xie Z, Yuan Y, et al. Plin5 alleviates myocardial ischaemia/reperfusion injury by reducing oxidative stress through inhibiting the lipolysis of lipid droplets. Sci Rep 2017;7:42574.
2. Farooq SM, Hou Y, Li H, et al. Disruption of GPR35 Exacerbates Dextran Sulfate Sodium-Induced Colitis in Mice. Dig Dis Sci 2018;63:2910-2922.
3. Li C, Roy K, Dandridge K, et al. Molecular assembly of cystic fibrosis transmembrane conductance regulator in plasma membrane. J Biol Chem 2004;279:24673-84.
4. Li C, Krishnamurthy PC, Penmatsa H, et al. Spatiotemporal coupling of cAMP transporter to CFTR chloride channel function in the gut epithelia. Cell 2007;131:940-51.
5. Lee CA, Chin LS, Li L. Hypertonia-linked protein Trak1 functions with mitofusins to promote mitochondrial tethering and fusion. Protein Cell 2018;9:693-716.
6. Kang T, Lu W, Xu W, et al. MicroRNA-27 (miR-27) targets prohibitin and impairs adipocyte differentiation and mitochondrial function in human adipose-derived stem cells. J Biol Chem 2013;288:34394-402.
7. Khadka M, Todor A, Maner-Smith KM, et al. The Effect of Anticoagulants, Temperature, and Time on the Human Plasma Metabolome and Lipidome from Healthy Donors as Determined by Liquid Chromatography-Mass Spectrometry. Biomolecules 2019;9.
8. Hsiao K, Zegzouti H, Goueli SA. Methyltransferase-Glo: a universal, bioluminescent and homogenous assay for monitoring all classes of methyltransferases. Epigenomics 2016;8:321-39.

### Supplementary Table 1. Primer sequences for genotyping

| Primers for Smyd5 genotyping | | | Note |
|---|---|---|---|
| Primer | Sequence 5’ → 3’ | Primer type | Designed by EMMA |
| Smyd5_F | GGTCTCATGGGGAACTGAGG | WT and mutant forward | |
| Smyd5_R | GCTTTCAGCCAAGCCAAGTC | WT reverse | |
| CAS_R1_Term | TCGTGGTATCGTTATGCGCC | Mutant Reverse | |
| Primers for Vil1-Cre genotyping | | | |
| 18960 | TTCTCCTCTAGGCTCGTCCA | Transgene Reverse | Designed by Jackson Lab |
| 14506 | CATGTCCATCAGGTTCTTGC | Transgene Forward | |
| oIMR7338 | CTAGGCCACAGAATTGAAAGATCT | Internal Positive control Forward | |
| oIMR7339 | GTAGGTGGAAATTCTAGCATCATCC | Internal Positive control Reverse | |

### Supplementary Table 2. Primer sequences for RT-qPCR

| Gene | Forward 5’ → 3’ | Reverse 5’ → 3’ |
|---|---|---|
| Tfam (human) | AAGATTCCAAGAAGCTAAGGGTGA | CAGAGTCAGACAGATTTTTTCCAGTTT |
| Cox I (human) | CCCACCGGCGTCAAAGTATT | TACAATGCCAGTCAGGCCAC |
| Gapdh (human) | ACCCACTCCTCCACCTTTGA | CTGTTGCTGTAGCCAAATTCGT |
| NADH dehydrogenase (human) | CGATTCCGCTACGACCAACT | GTTTGAGGGGGAATGCTGGA |
| GAPDH (human) | TTTCTTTGCAGCAATGCCTCC | CCATTCCCCAGCTCTCATACC |
| Cox I (mouse) | GGTCAACCAGGTGCACTTTT | TGGGGCTCCGATTATTAGTG |
| Cox II (mouse) | CCACTTCAAGGGAGTCTGGA | AGTCATCTGCTACGGGAGGA |
| Gapdh (mouse) | CATCGTGGAAGGGCTCATGAC | CTTGGCAGCACCAGTGGATG |
| Pgc1a (mouse) | TGAATGCAGCGGTCTTAGCA | TGCTCCATGAATTCTCGGTCTTA |

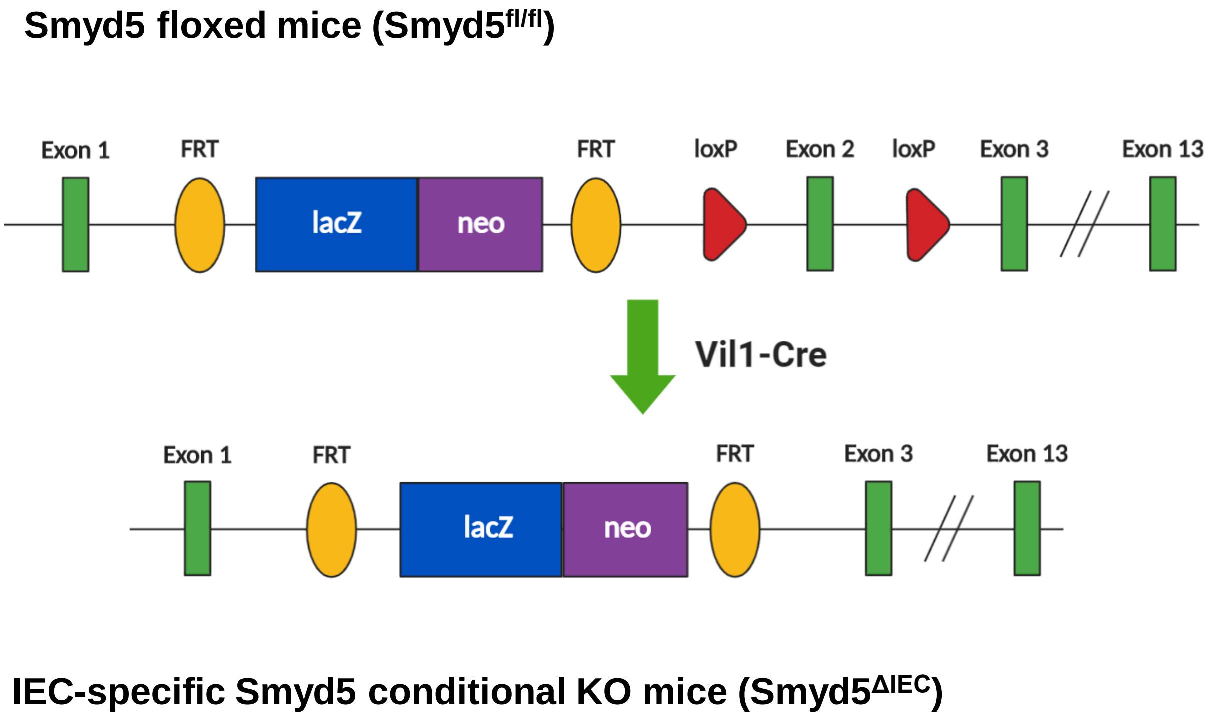


**Supplementary Figure 1. Generation of mice with a conditional null allele for Smyd5.**

Smyd5 allele containing LacZ-reporter-promoter (blue box)-driven neo targeting cassette (purple box), FLP-FRT sites (yellow oval), Cre-loxP sites (Red triangle), and Smyd5 exons (green rectangle). Breeding of Smyd5 floxed mice (Smyd5$^{fl/fl}$) with Vil-Cre transgenic mice resulted in truncated Smyd5 copy (with exon 2 deleted), denoted IEC-specific Smyd5 depletion (Smyd5$^{\Delta IEC}$).

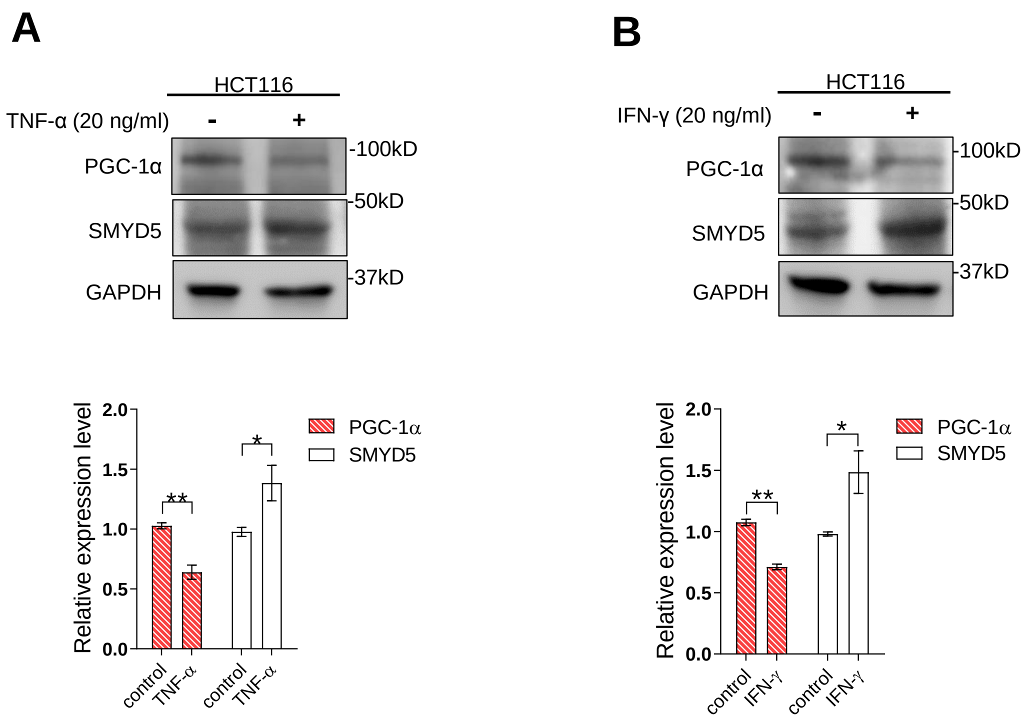


**Supplementary Figure 2. SMYD5 is up-regulated and PGC-1α is down-regulated in HCT116 cells upon the treatment of proinflammatory cytokines.**

HCT116 cells were treated with (**A**) TNF-α (20 ng/ml) or (**B**) IFN-γ (20 ng/ml) for 24 hrs, and the expression levels of SMYD5 and PGC-1α were determined by Western blot analysis. The quantitative analyses of the band intensity of SMYD5 and PGC-1α in TNF-α or IFN-γ treated HCT116 cells were also presented. *$p$ < 0.05, **$p$ < 0.01; n = 3.

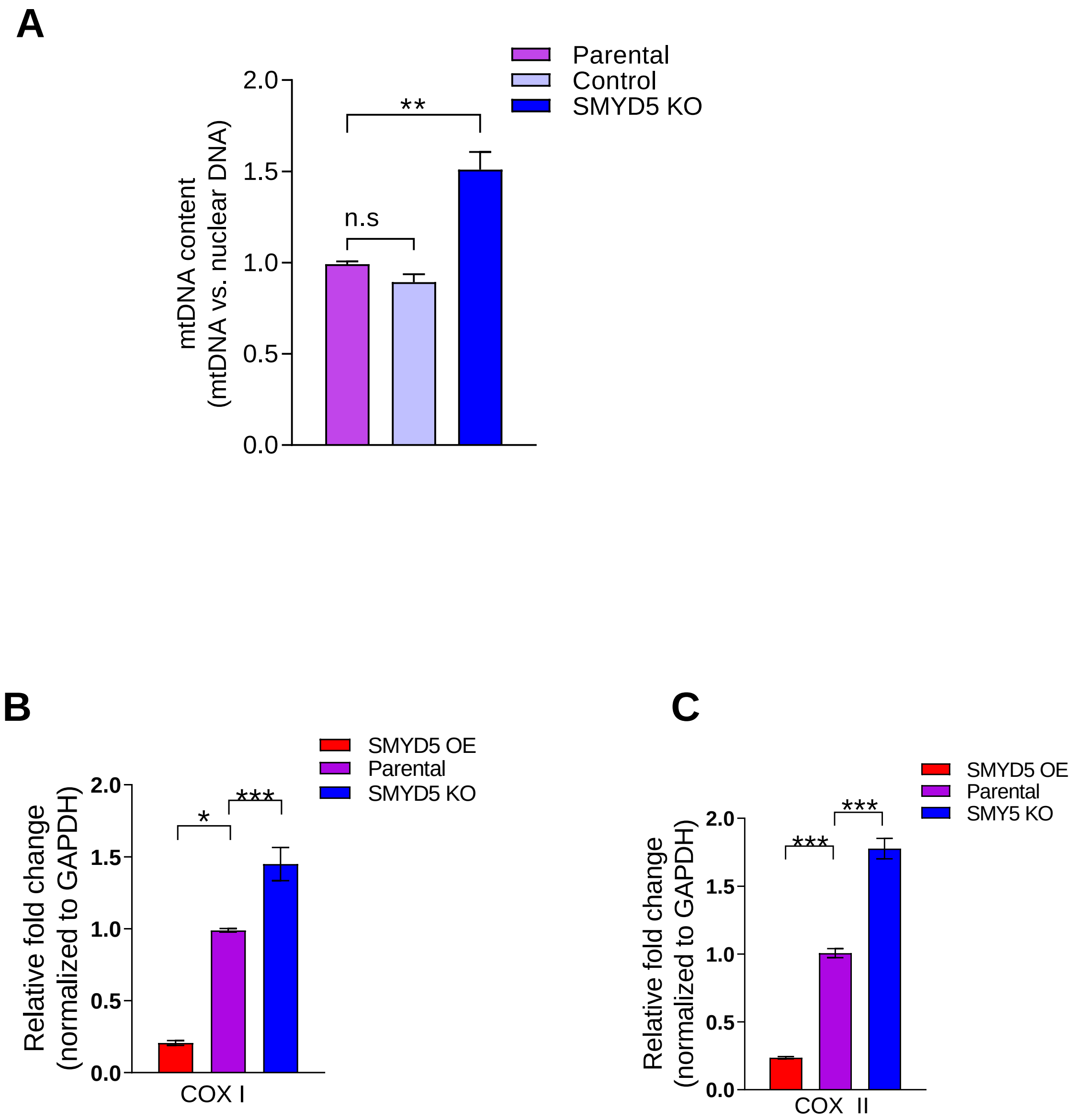


**Supplementary Figure 3. SMYD5 regulates mitochondrial biogenesis in HCT116 cells.**

RT-qPCR analysis of the abundance of mitochondrial DNA (mtDNA) content (**A**) and the expression of the mtDNA-encoded genes, the mitochondrial biogenesis markers COX I (**B**) and COX II (**C**) in HCT116 cells with overexpressing (OE) or knockout (KO) SMYD5.
The relative mtDNA content was determined using ΔCT method. The primers for human mtDNA-encoded Cox II gene and for human nuclear DNA-encoded GAPDH were used for the analysis. The results were presented as the ratio of mtDNA relative to nuclear DNA.
Results were presented as fold changes relative to parental HCT116 cells. *$p$ < 0.05, **$p$ < 0.01, ***$p$ < 0.001; n.s., no significance; n = 3-4.

**A**

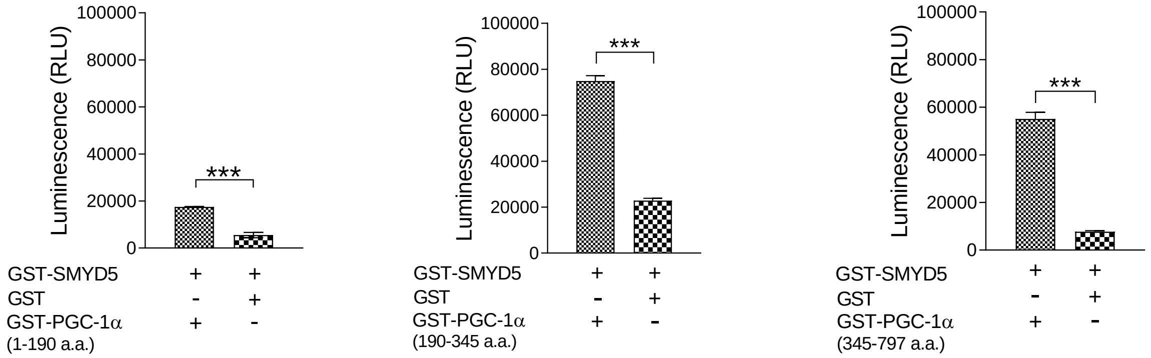


**B**

### Fragment Ion Calculator Results

Sequence: **YLTTNDDPPHTKPTENR**, pI: **5.38094**

#### Fragment Ion Table, monoisotopic masses

| Seq | # | B | Y | # (+1) |
|---|---|---|---|---|
| Y | 1 | 164.07065 | 2012.96670 | 17 |
| L | 2 | 277.15471 | 1849.90337 | 16 |
| T | 3 | 378.20239 | 1736.81931 | 15 |
| T | 4 | 479.25007 | 1635.77163 | 14 |
| N | 5 | 593.29300 | 1534.72395 | 13 |
| D | 6 | 708.31994 | 1420.68102 | 12 |
| D | 7 | 823.34688 | 1305.65408 | 11 |
| P | 8 | 920.39965 | 1190.62714 | 10 |
| P | 9 | 1017.45241 | 1093.57437 | 9 |
| H | 10 | 1154.51132 | 996.52161 | 8 |
| T | 11 | 1255.55900 | 859.46270 | 7 |
| K | 12 | 1397.66906 | 758.41502 | 6 |
| P | 13 | 1494.72183 | 616.30496 | 5 |
| T | 14 | 1595.76950 | 519.25219 | 4 |
| E | 15 | 1724.81210 | 418.20452 | 3 |
| N | 16 | 1838.85502 | 289.16192 | 2 |
| R | 17 | 1994.95613 | 175.11900 | 1 |

#### Mass/Charge Table

| | Mass | |
|---|---|---|
| | Mono | Avg |
| (M) | 2011.95942 | 2013.13919 |
| $(M+H)^{+}$ | 2012.96670 | 2014.14646 |
| $(M+2H)^{2+}$ | 1006.98701 | 1007.57689 |
| $(M+3H)^{3+}$ | 671.66045 | 672.05370 |
| $(M+4H)^{4+}$ | 503.99717 | 504.29211 |

Modifications:
* To residue 12 added the value 14.0151

Sequences in Protein:
K.YLTTNDDPPHT**K**PTENR.N

PSM Modification Positions in Protein:
1×Methyl [**K223**]

Confidence: High

**Supplementary Figure 4. PGC-1α is mono-methylated at lysine 223 by SMYD5.**
(**A**) Luminescence-based in vitro methyltransferase assay of PGC-1α methylation mediated by SMYD5. Bacterially purified GST-SMYD5 was mixed with GST alone or GST-PGC-1α fragments (GST-PGC-1α aa 1-190, GST-PGC-1α aa 190-345, and GST-PGC-1α aa 345-797) in an assay reaction that detects S-adenosyl-L-homocysteine, the universal reaction products of all methyltransferases, as described in the Methods. (**B**) LC-MS/MS analysis showed methylation of PGC-1α at lysine 223. Theoretical values of MS fragments are summarized. ***$p < 0.001$; n = 3.

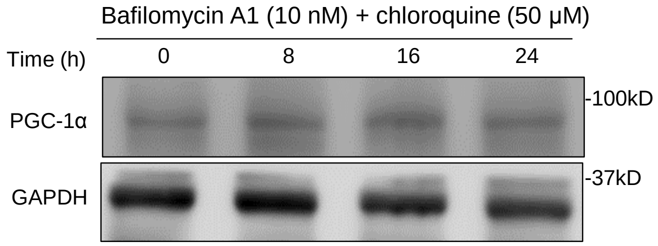


**Supplementary Figure 5. Lysosomal inhibition does not affect the protein level of PGC-1α in HCT116 cells.**

HCT116 cells were treated with a combination of inhibitors of autophagy-lysosomal pathway: bafilomycin A1 (10 nM) and chloroquine (50 μM), for 0, 8, 16, and 24 hrs, respectively. Whole cell lysates were immunoblotted for the expression of PGC-1α and GAPDH using respective antibodies.

A

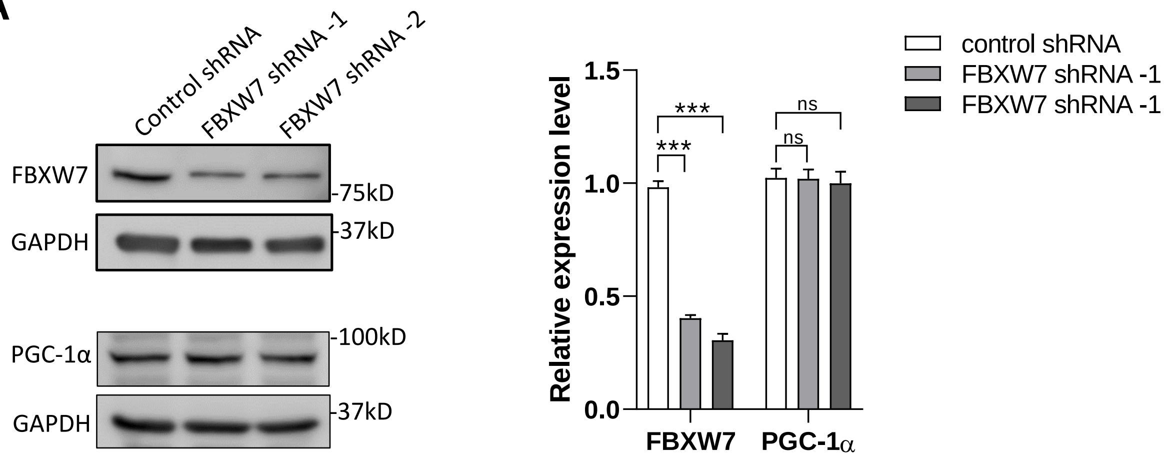


B

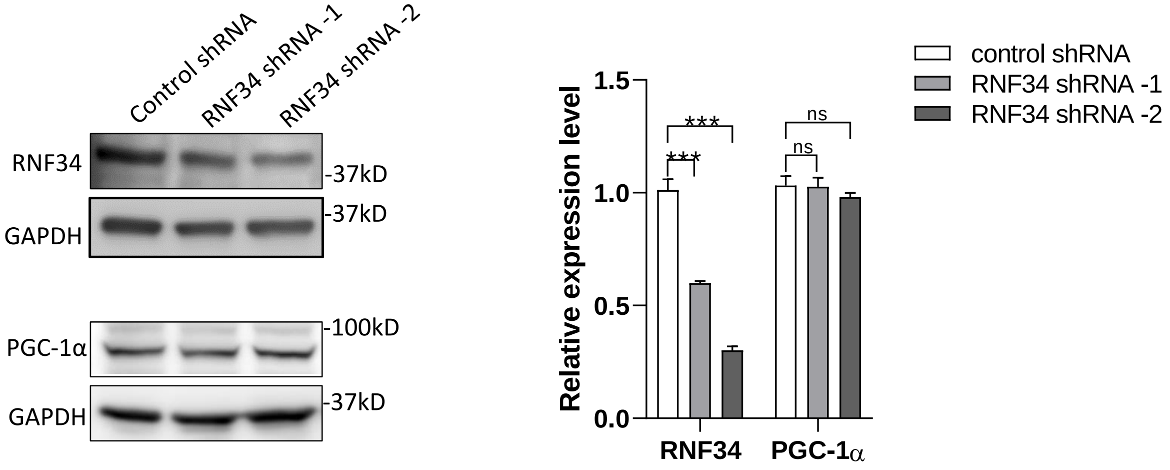


**Supplementary Figure 6. Depletion of FBXW7 or RNF34 does not affect the expression of PGC-1α in IECs.**

(**A**) HCT116 cells were transduced with shRNA viruses targeting the E3 ligase FBXW7 (FBXW7 shRNA -1 or -2) or control shRNA viruses, and whole cell lysates were immunoblotted for FBXW7, PGC-1α, and GAPDH. $^{***}p < 0.001$ (n = 3); ns, not significant.

(**B**) HCT116 cells were transduced with shRNA viruses targeting the E3 ligase RNF34 (RNF34 shRNA -1 or -2) or control shRNA viruses, and whole cell lysates were immunoblotted for RNF34, PGC-1α, and GAPDH. $^{***}p < 0.001$ (n = 3); ns, not significant.

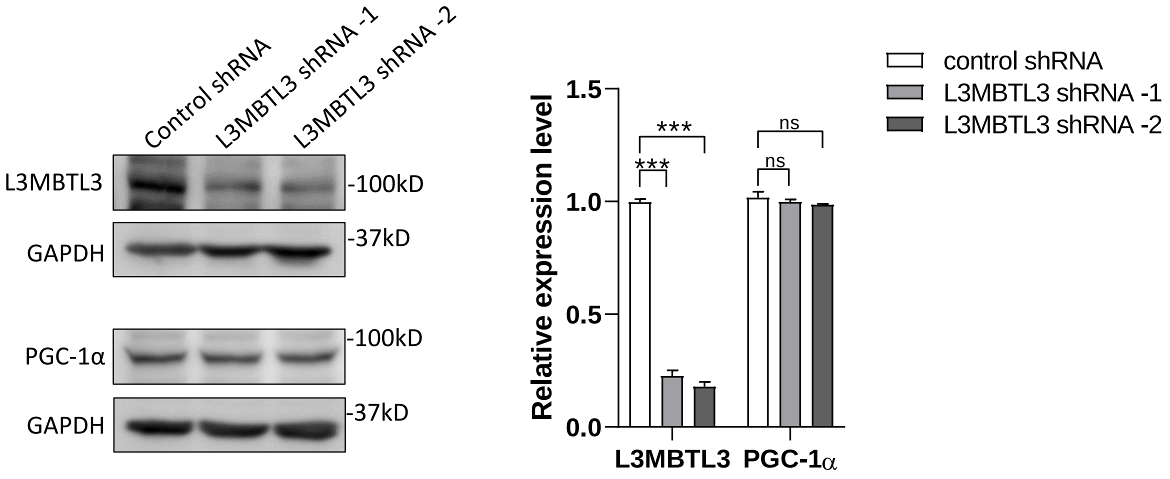


**Supplementary Figure 7. Depletion of L3MBTL3 does not affect the expression of PGC-1α in IECs.**

HCT116 cells were transduced with shRNA viruses targeting the methyl-binding protein L3MBTL3 (L3MBTL3 shRNA -1 or -2) or control shRNA viruses, and whole cell lysates were immunoblotted for L3MBTL3, PGC-1α, and GAPDH. $^{***}p < 0.001$ (n = 3); ns, not significant.
